# Comparative genomic analysis of *Planctomycetota* potential towards complex polysaccharide degradation identifies phylogenetically distinct groups of biotechnologically relevant microbes

**DOI:** 10.1101/2024.01.10.575047

**Authors:** Dominika Klimek, Malte Herold, Magdalena Calusinska

## Abstract

The outstanding hydrolytic potential of the *Planctomycetota* phylum for complex polysaccharide degradation has recently been acknowledged based on the numerous carbohydrate-active enzymes (CAZymes) encoded in their genomes. However, mainly members of the *Planctomycetia* class have been characterised up to now, and little is known about the degrading capacities of the other *Planctomycetota*. Our in-depth characterisation of the available planctomycetotal genomic resources increased our knowledge of the carbohydrolytic capacities of *Planctomycetota*. We showed that this single phylum encompasses a wide variety of the currently known CAZyme diversity assigned to glycoside hydrolase families, and that many members are characterised by a high versatility towards complex carbohydrate degradation, including lignocellulose. We also highlighted members of the *Isosphaerales, Pirellulales, Sedimentisphaerales* and *Tepidisphaerales* orders as having the highest encoded hydrolytic potential of the *Planctomycetota*. Furthermore, members of a yet uncultivated group affiliated to *Phycisphaerales* were identified as an interesting source of novel, lytic polysaccharide monooxygenases that could boost lignocellulose degradation. Surprisingly, many *Planctomycetota* from anaerobic digestion reactors were shown to encode CAZymes targeting algal polysaccharides – this opens new perspectives for algal biomass valorisation in biogas processes. Our study provides a new perspective on planctomycetotal carbohydrolytic potential, highlighting distinct phylogenetic groups which could provide a wealth of diverse, potentially novel CAZymes of industrial interest.

## Introduction

Modern society generates enormous amount of waste organic matter that requires specific and well-defined disposal procedures [1]. Instead, waste biomass could be valorised into added-value products and energy [2]. In the light of the global threat related to the environmental pollution and climate change, the bioconversion of organic waste into biofuels and sustainable, added-value products has been gaining considerable attention. In nature, the degradation of organic matter is mediated by both aerobic and anaerobic microorganisms capable of producing the broad assortment of hydrolytic enzymes [3]. The enzymes involved in carbohydrate metabolism are known as carbohydrate-active enzymes (CAZymes), and they are classified into four classes that include glycoside hydrolases (GH), carbohydrate esterases (CE), polysaccharide lyases (PL) and enzymes with auxiliary activities (AA) [4]. Carbohydrate binding modules (CBMs) are non-catalytic modules, generally defined as accessory CAZymes, and their main role is to recognise the substrates by binding carbohydrates [5]. Several different industrial sectors, such as food industries and biorefineries, have been benefiting from various bacterial and fungal strains for CAZymes provision [6]. A better understanding of microbial physiology and ecology could result in enlarging the pool of available strains for scientific and industrial use, such as recombinant design and enzyme cocktail development [7].

Although a number of metagenomic studies have revealed a high diversity of complex polysaccharide-degrading microbes in distinct biomass-rich habitats, little attention has been paid to *Planctomycetota* [3, 8, 9]. According to a recent analysis of a global distribution of carbohydrate utilisation potential in the tree of life, alongside *Bacteroidota, Fibrobacterota, Acidobacteriota*, and a few other phyla, *Planctomycetota* was identified as a phylum whose members are the most versatile in degrading diverse biopolymers of cellulosic and non-cellulosic origin [10]. *Planctomycetota*, previously known as *Planctomycetes*, is one of the phyla within the *Planctomycetota-Verrucomicrobiota-Chlamydiota* superphylum (PVC). They are characterised with distinctive features not commonly detected in other prokaryotes, such as enlarged periplasm, outer membrane complexes in the form of crateriform structures, and a non-FtsZ based division mode [11, 12]. Besides these cellular particularities, bacteria belonging to this widespread phylum have been highlighted in different environments for their hydrolytic potential [13]. Uncultured *Planctomycetota* have been identified as primary degraders of extracellular polymeric substances in soil [14] and complex carbohydrates in marine sediments [15, 16]. Certain members of this phylum can attach to algal surfaces and were proposed to depolymerise the algal-derived polymers [17, 18]. *Rhodopirellula* is widely known for producing multiple and diverse sulphatases engaged in sulphated polysaccharides degradation [19]. Metatranscriptomic studies also revealed the contribution of *Planctomycetota* to complex polysaccharide degradation in *Sphagnum*-dominated areas [20]. Additionally, their cellulo-, hemicellulo-, and chitinolytic activities have been reported in environments like wetland and soil, predominantly inhabited by the *Gemmatales* and *Isosphaerales* orders of the *Planctomycetia* class [13, 21]. However, so far, only a limited number of *Planctomycetota* have been considered in applied settings [22], and most of the characterised strains derive from the *Planctomycetia* class only [23].

In this study we aimed to expand the knowledge on the carbohydrate-degrading capacities of *Planctomycetota*, including the *Planctomycetia* and *Phycisphaerae* classes, as well as other less characterised members of this phylum. To do so, we employed different bioinformatic approaches to the collection of 1425 non-redundant planctomycetotal genomes retrieved from own and public repositories. As a result, we show that a high number of diverse CAZymes are encoded in different orders of *Planctomycetota*, indicating the wide versatility of the different members towards carbohydrates degradation. At the same time, a relatively low number of CAZyme families are shared between the *Planctomycetota* orders, further indicating within-phylum specialisation of the different planctomycetotal groups towards different fractions of polysaccharides constituting biomass. High incidence of CAZyme gene clusters and presence of potentially extracellular enzymes, point to the existence of coordinated microbial strategies in complex polysaccharides degradation, including lignocellulose and algal biomasses.

## Methods

### Dataset acquisition and classification of planctomycetotal genomes

Over 3000 publicly available draft and complete planctomycetotal genomes were downloaded from GenBank in June 2021 (http://www.ncbi.nlm.nih.gov/genbank/), using ncbi-genome-download (https://github.com/kblin/ncbi-genome-download). The initial genome database was complemented with the metagenome-assembled genomes (MAGs) from own studies (see Additional File 1, Table S1 for further details) as well as genomes from the catalogue of Earth’s Microbiomes (GEM) available from the JGI [24]. Unless stated otherwise, we will refer to both individual genomes and MAGs as genomes. All genomes were assessed for redundancy using dRep v3.2.2 with the -con 10 --checkM_method taxonomy_wf parameters [25]. The resulting 1457 non-redundant genomes were classified taxonomically with GTDB-tk v1.2.0 against GTDB database release 89 [26] and only hits designated as *Planctomycetota* were retained. A few changes were introduced to the classification retrieved. Briefly: taxonomic names were refined by removing prefixes, the *Planctomycetes* class was renamed *Planctomycetia* and the candidate order UBA1161 was renamed *Tepidisphaerales* according to Dedysh et al. [27]. CheckM v1.2.0 was used to determine genome completeness and contamination [28] and only genomes meeting the MIMAG standard of medium to high quality level (i.e. completeness above 75% and contamination below 10%) were retained for further analyses [29]. At this stage, our database contained 1451 non-redundant planctomycetotal genomes. We further reduced this number to 1425, by excluding the genomes encoding less than 10 carbohydrate active enzymes (CAZymes). To simplify the analysis, we designated classes with fewer than 10 sequenced genomes as “Other class”. The final database of 1425 non-redundant genomes was additionally complemented with the respective metadata retrieved from NCBI using the rentrez script [30], followed by manual curation for conflicting information. Environmental metadata from the GEM catalogue was retrieved directly from the deposited repository and unified with the NCBI entries for the different habitat categories (Additional File 1, Table S2).

### Functional annotation of planctomycetotal genomes

Genomes were gene-called by prodigal v2.6.3 [31] and CAZymes were annotated by dbCAN2 v2.0.11 [32] against the dbCAN database v9 [4] using the three integrated tools (DIAMOND, Hotpep and, HMMER) with default parameters [33–35]. Genome annotation was also performed using Prokka v1.14.6 with its default databases [36]. To determine clusters of co-localized CAZymes, we applied a modified version of the CGCFinder module of dbCAN2 [32] to detect CAZyme gene clusters (CGCs). CGCs were predicted as gene clusters consisting of at least one CAZyme coding gene with at least one auxiliary gene (*e.g.* transcription factors or transporters) or another CAZyme separated by at most two other genes. Positive hits were assigned to CAZymes families if annotated by HMMER v3.1.2 and multiple CAZyme assignments were considered as separate functional domains or modules. For searching putatively novel CAZymes only hits annotated either by DIAMOND v0.9.19 or Hotpep, but not HMMER, were retained. To assess the novelty of predicted CAZymes (assigned by HMMER), we searched the protein sequences against the CAZy database with DIAMOND and amino acid sequence identity of the best hit was inferred. Signal peptides were detected using signalP v6 [37]. Glycosyltransferase coding genes were excluded from the analysis as they are not involved in polysaccharide degradation.

### Data analysis

Statistical analyses and visualisations were performed with R software v 4.0.2 [38]. For multivariate analyses, a presence-absence table of CAZyme content for each genome was transformed into a Jaccard distance matrix (Additional File 2, Table S3). CAZyme dissimilarity was assessed using principal coordinate analysis (PCoA) and permutational ANOVA (PERMANOVA) as well as analysis of similarities (ANOSIM) with the vegan v 2.5.7 package in R [39]. Linear discriminant analysis (LDA) was performed with a nonparametric Kruskal Wallis test using the microbial v0.0.22 package in R (logarithmic LDA score > 4) [40]. Clustering genomes with similar set of CAZymes was assessed by Dirichlet multinomial mixtures method using mothur v1.48.0 [41]. A phylogenetic tree of genomes was constructed from the alignment of default marker genes using PhyloPhlAn v3.0.60 (--diversity medium supertree_aa) [42]. The alignment of protein sequences was calculated using the MUSCLE algorithm with its default parameters [43]. Pairwise comparisons between protein sequences and Neighbor-Joining consensus was calculated for constructing the tree using Geneious Prime v 2019.0.3 [44]. The Spearman’s rank correlation was calculated in R using package stats. Unless otherwise stated, the significance of differences between tested groups was assessed using either a non-parametric Kruskal-Wallis or Wilcoxon test (R package stats). The obtained p-values were adjusted for multiple testing using the Benjamini–Hochberg procedure (false-discovery rate).

### Annotation of CAZyme family activities

The substrate database was framed according to [10] and the CAZy database [4] (Additional File 3). For CAZyme functional analysis, entries assigned to GH and PL families were classified based on their main characterised enzymatic activities into four categories according to the main target: algal biomass (algal-derived polysaccharides), plant biomass (plant storage polysaccharides, oligosaccharides, and cellulose-hemicellulose fractions), algal/plant biomass and other activities (all the remaining polysaccharide targets were grouped together). Further, the categories were subdivided based on the substrate specificity: algal polysaccharides, glucans (α- and β-glucans), oligosaccharides, lignocellulose (cellulosic and/or hemicellulosic polysaccharides), NAG-based polysaccharides (based on N-acetylglucosamine, NAG), pectin, and other polysaccharides. The detailed annotations of substrates are available in the Additional File 3, Table S1. The ratio of CAZymes for polysaccharide target specificity was calculated by comparing the number of CAZymes with assigned function to the number of all predicted CAZymes.

## Results

### Database of planctomycetotal genomes

As most of the *Planctomycetota* remain uncultured, we created a database of 1425 non-redundant and medium to high quality genomes (see Methods), recovered from both metagenomics and isolate sequencing studies (Additional File 1, Table S1). Specifically, the database included 662 genomes of the *Planctomycetia* class with the following orders: *Gemmatales* (number of genomes, n=87)*, Isosphaerales* (n=28)*, Pirellulales* (n=408), and *Planctomycetales* (n=137). Furthermore, the database included 463 genomes of the *Phycisphaerae* class including the *Phycisphaerales* (n=246)*, Sedimentisphaerales* (n=118), and *Tepidisphaerales* (n=13), as well as putative UBA1845 (n=64) and SM23-33 (n=22) orders (Additional File 1, Table S1). *Planctomycetia* and *Phycisphaerae* are the two biggest and widely described classes of the *Planctomycetota* phylum, and a few isolated strains of these classes are the only so far cultured and characterised carbohydrate degraders [12]. Another assigned class was much less numerous and included 46 genomes of the *Brocadiae* candidate class, which are commonly known as anaerobic ammonium oxidising (anammox) bacteria widely employed in wastewater treatment settings [45]. Other genomes (172) represented novel, not yet assigned planctomycetotal classes, including UBA8742, UBA8108, UBA1135 and UBA11346, which we labelled under a cluster name of “putatively novel classes” (Fig. 1a). Genomes that represent other less populated classes of *Planctomycetota* (<10 genomes) were grouped together as “Other” class (see Methods, n=65).

**Fig. 1.**
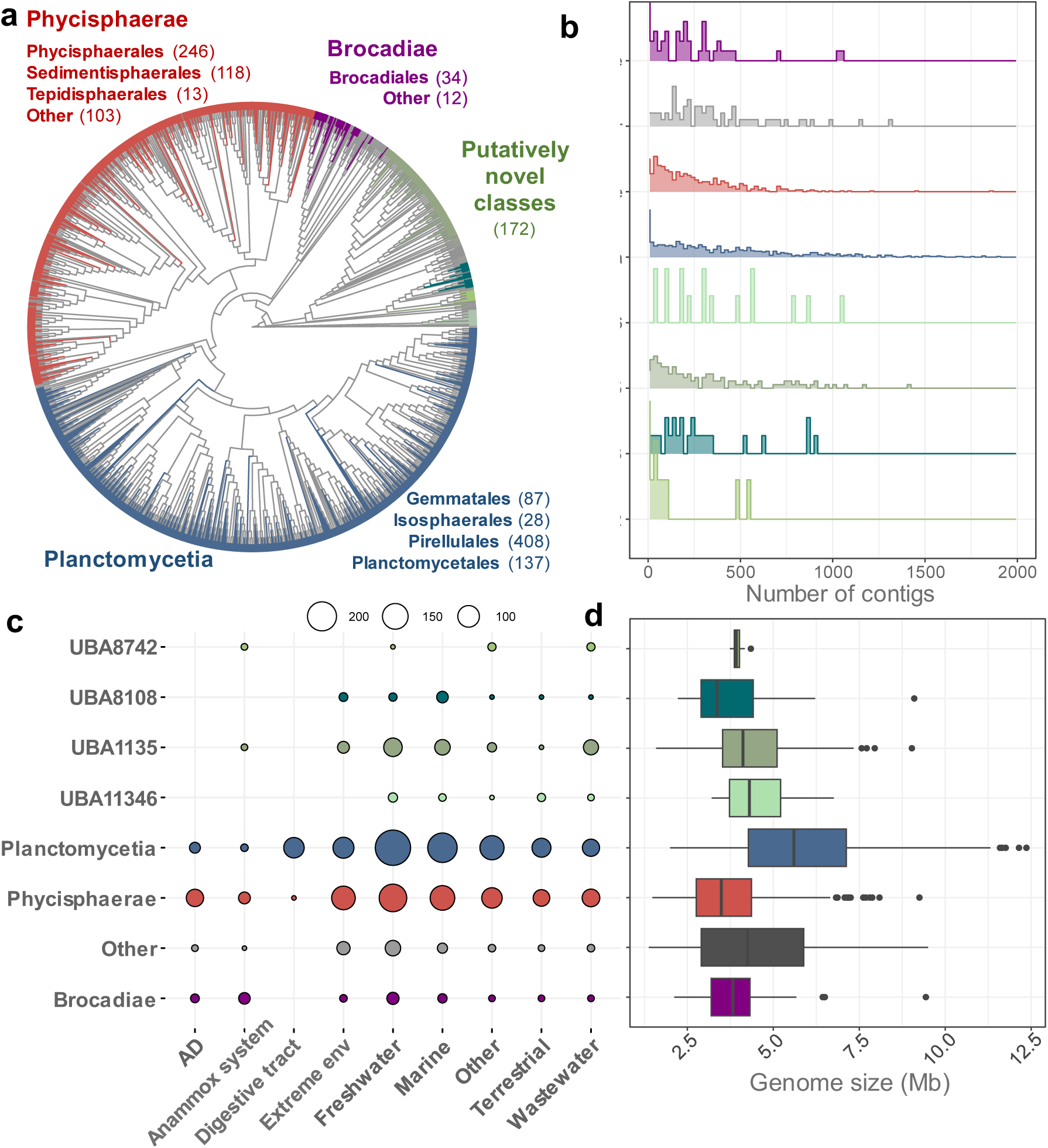
Overview of planctomycetotal genomes included in the study, grouped and coloured at the class level **a.** The phylogenetic distribution of planctomycetotal genomes. The grey colour on the outer circle represents genomes excluded from further analysis (see methods section for details) **b.** Number of contigs in genome assemblies **c-d.** Environmental origin, further called “habitat” (c) and genome size in Mb (d) of planctomycetotal genomes. Circle size corresponds to the number of genomes, according to the legend on the top.

The majority of planctomycetotal genomes included in our database consists of less than 250 contigs (Fig. 1b). Nevertheless, the genomes of the *Planctomycetia* class are in many cases highly fragmented, but their members are characterised with the greatest genome sizes among all the *Planctomycetota* phylum (Fig. 1d). For instance, phycisphaeral and brocadial genomes have smaller range sizes (3.6 – 3.9 Mb on average), thus being easier to reconstruct. According to the environmental metadata, half of the planctomycetotal genomes in our database originated from marine and freshwater habitats (51%) while the remaining genomes were retrieved from extreme environments including thermal springs, hydrothermal vents and saline/alkaline habitats (13%), wastewater (8%), terrestrial (7%), animal digestive systems (4%), anammox (2%), AD reactors (4%) and other environments (11%) (Fig. 1c; Additional File 1, Table S2).

Our database demonstrates virtually all the currently known planctomycetotal diversity based on the position in the phylogenetic tree (Fig. 1a) and a broad spectrum of environments where *Planctomycetota* naturally occur [23], highlighting differences between groups. This comprehensive collection allowed us to largely complement another recent study of microbial CAZymes, which included only 243 planctomycetotal genomes [10].

### Within-phylum and within-class distribution of planctomycetotal CAZymes

The size of the CAZyme repertoire (CAZyome) in bacterial genomes is directly correlated with a higher potential for polysaccharide degradation [46]. Therefore, we assessed the set of encoded CAZyme families in planctomycetotal genomes retained in our database to estimate their catalytic potential. Globally, we detected as many as 232 CAZyme families and 132 CAZyme subfamilies (Additional File 2, Table S3). At the time of writing (November 2023), 187 GH families are listed in the CAZy database [4]. Here, 138 different GH families were detected in the analysed planctomycetotal genomes, which demonstrates that this single phylum covers 80% of the known GH family diversity at the time of analysis (Additional File 2, Table S3). In turn, the diversity of planctomycetotal AAs, CEs, PLs and CBMs represent respectively 53%, 70%, 69% and 43%, of the described CAZyme classes (family diversity). Knowing that AAs and PLs compared to GHs and CEs are much less occurring between bacterial phyla [10], these numbers are quite high for a single phylum.

First, we calculated the most and the least encoded CAZyme families, employing respective thresholds: present in at least 50% and present in less than 5% of analysed planctomycetotal genomes. As a result, we identified 17 commonly occurring CAZyme families (i.e. encoded by >50% of genomes), including six GHs, four CBMs and seven CEs (Additional File 2, Table S4). The most encoded GHs included GH2, GH5, GH13, GH33, GH57 and GH109, all of which can be largely classified as broad spectrum oligosaccharidases, and α/β-glucanases (Additional File 3, Table S1). Subsequently, we verified the most frequently encoded CAZyme families within specific planctomycetotal classes i.e. >50% of representatives assigned to the same class (Additional File 7, Fig. S1). Almost all of the genomes affiliated to the *Brocadiae* candidate class (mainly *Brocadiaceae*) encode the key α-glucanases such as GH13, GH15, GH57, GH77, suggesting their capacity to utilise α-1,4- and α-1,6-glucans such as starch or glycogen. Members of UBA8742 have a very limited number of CAZyme families, but a high fraction of genomes encodes specific CAZymes putatively engaged in hemicelluloses (GH44) or pectin (PL9) degradation (Additional File 7, Fig. S1). Genomes belonging to the *Planctomycetia* class are deprived of members encoding GH102 family, which are present in all other planctomycetotal classes. Intriguingly enough, some genomes of *Planctomycetota* encode CAZyme coding genes assigned to AA12 family (Additional File 2, Table S3) which has never been detected in any prokaryote so far [4]. Nevertheless, these observations would require further investigation in order to clarify their relevance. In turn, when looking at the least encoded CAZymes (i.e. encoded by <5% of genomes), 110 CAZy families were identified, including multiple GHs and CBM domains (Additional File 2, Table S5). We could notice that many of these CAZymes were frequently encountered in the *Phycisphaerales* (the *Phycisphaerae* class) and *Planctomycetales* orders (the *Planctomycetia* class), whose genomes encode 52% and 44% of rare CAZyme families respectively.

To assess how uniformly the different CAZymes are distributed in the genomes of *Planctomycetota*, we looked at the number of all the CAZyme families that can be found within each planctomycetotal class (Fig. 2). We verified that 129 CAZyme families are shared by all the planctomycetotal classes while only 6 CAZyme families are unique to the different *Planctomycetota* classes (Fig. 2a). The *Phycisphaerae* class encodes the highest number of CAZyme families (14) not found in any other planctomycetotal class such as β-agarases GH118, mannan-targeting GH47 and GH134 or xylan-targeting GH11 and GH54 (Fig. 2a). The *Planctomycetia* class shares with *Phycisphaerae* 20 CAZyme families including endo-glucanases such as GH12 and GH64, GH98 endo-xylanases or PL4 rhamnogalacturonan lyses (Fig. 2a).

**Fig. 2.**
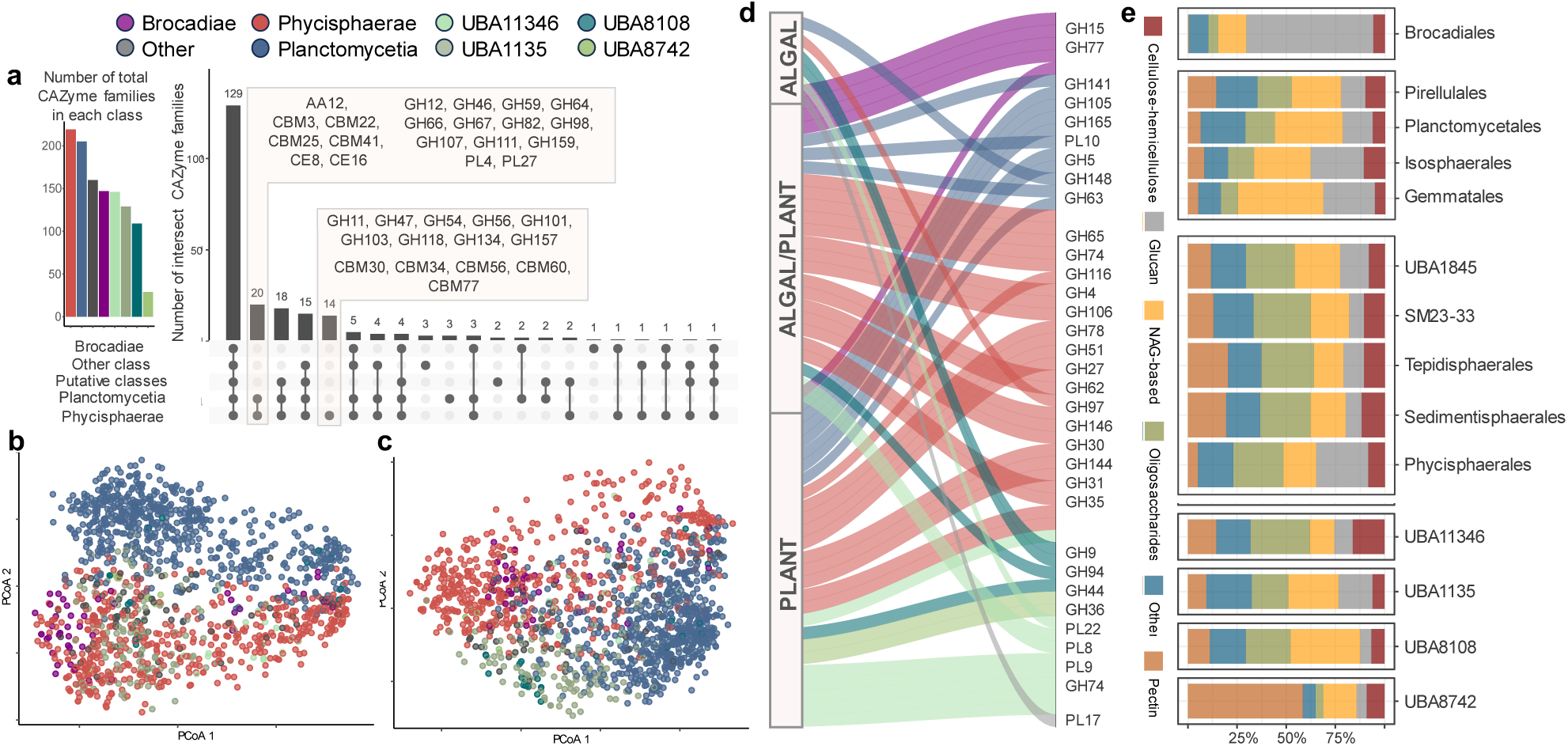
Characterisation of planctomycetotal CAZyomes (CAZyme repertoires) coloured by class affiliation; only main trends are shown. **a.** Bar plot (left) and Upset (right) plot representing the number of CAZyme families and intersections between the planctomycetotal classes. **b-c.** Principal coordinates analysis (PCoA) ordination based on the Jaccard distance presence-absence matrix of GHs (b) and all the other CAZyme families (c) encoded in planctomycetotal genomes. **d.** Alluvial plot shows the number of significantly enriched GH and PL families between orders (p < 0.05) with assigned functions towards either type of biomass. **e.** Bar plot presenting functional assignment of GHs and PLs, coloured by the substrate category.

To compare within-class planctomycetal CAZyomes, we employed the principal coordinate analysis (PCoA, Fig. 2b-c) applied to the CAZyme occurrence matrix. We revealed a moderate separation between the different planctomycetotal classes, especially visible for GH families (Fig. 2b), which was further supported by statistical tests (PERMANOVA p < 0.01 and ANOSIM R=0.45 p < 0.01). Subsequently, we verified to what extent the lifestyle which is directed by the habitat specificity could also justify the observed separation of the planctomycetotal carbohydrate degradation potentials. Nevertheless, we found only a low impact of genome origin (ANOSIM R=0.06 p < 0.01), except a clear grouping for planctomycetotal genomes retrieved from animal digestive tract. Even though different genomes are sourced from very distantly related animals, such as insects (e.g. termites) and herbivorous mammals (e.g. camels, goats or pigs), all belong to the *Thermoguttaceae* family of the *Pirellulales* class (Additional File 1, Table S1).

### Phylogenetically distinct groups of *Planctomycetota* display redundant hydrolytic potentials

To specifically look for the different encoded CAZyme families in planctomycetotal genomes, we detected a panel of 101 differentially encoded CAZymes between the orders of *Planctomycetota* (Fig. 2d; Additional File 2, Table S6). Members of *Phycisphaerae* show the highest number of differentially encoded CAZymes targeting plant biomass (Fig. 2d), including hemicellulases GH10 (UBA1845 putative order), lyases of pectic polysaccharides PL4 and GH143 (*Tepidisphaerales* order), GH11 and GH115 targeting xylans (*Sedimentisphaerales* order). Multiple differentially encoded CAZymes among *Phycisphaerae* are predicted as algalytic e.g., β-agarases GH50, laminarinases GH128 and GH157 (*Tepidisphaerales* order), α-L-fucosidases GH151 and carrageenan sulphatases GH167 (SM23-33 order). Among the *Planctomycetia* class, members of *Pirellulales* order preferentially encode β-agarases GH86 or endo-xylanases/α-fucosidases GH141 while *Isosphaerales* order encode endo-glucanases/laminarinases GH64, endo-glucanases/lichenases GH148 as well as multi-functional GH5 family.

On one hand, the separation of CAZyomes between the main planctomycetotal classes, together with a high number of differentially enriched CAZymes could point towards distinct carbohydrate metabolisms of *Planctomycetota* members. On the other hand, it is widely known that different GH families may catalyse the hydrolysis of structurally similar substrates, as such diverse CAZyomes could be functionally redundant [47]. Therefore, to further assess the planctomycetotal carbohydrate degradation potentials, we broadly classified the differentially enriched CAZymes in planctomycetotal genomes to one of the biomass and substrate categories (Fig. 2e; see Methods). Overall, while different CAZyme families are preferentially encoded in different groups, all *Planctomycetota* seem equally well equipped to target all main biomass categories, independently of their phylogenetic origin and habitat specificity (Fig. 2e). Interestingly, genomes of marine bacteria are also enriched in lignocellulose degrading CAZymes, even though marine polysaccharides are different from terrestrial carbohydrates, and are often highly sulphated [48]. Conversely, planctomycetotal genomes retrieved from diverse environments such as freshwater or engineered systems (anammox and AD reactors) encode similar potential towards algal-derived polysaccharides (6.49% ± 0.58 algalytic CAZymes) than average marine *Planctomycetota* (Fig. 3f), suggesting they are equally well-suited to targeting algal biomass. Simultaneously, AD-originated *Planctomycetota* are significantly enriched (p < 0.01) in enzymes specific to diverse sulphated N-glycans, especially fucose-based polysaccharides.

**Fig. 3.**
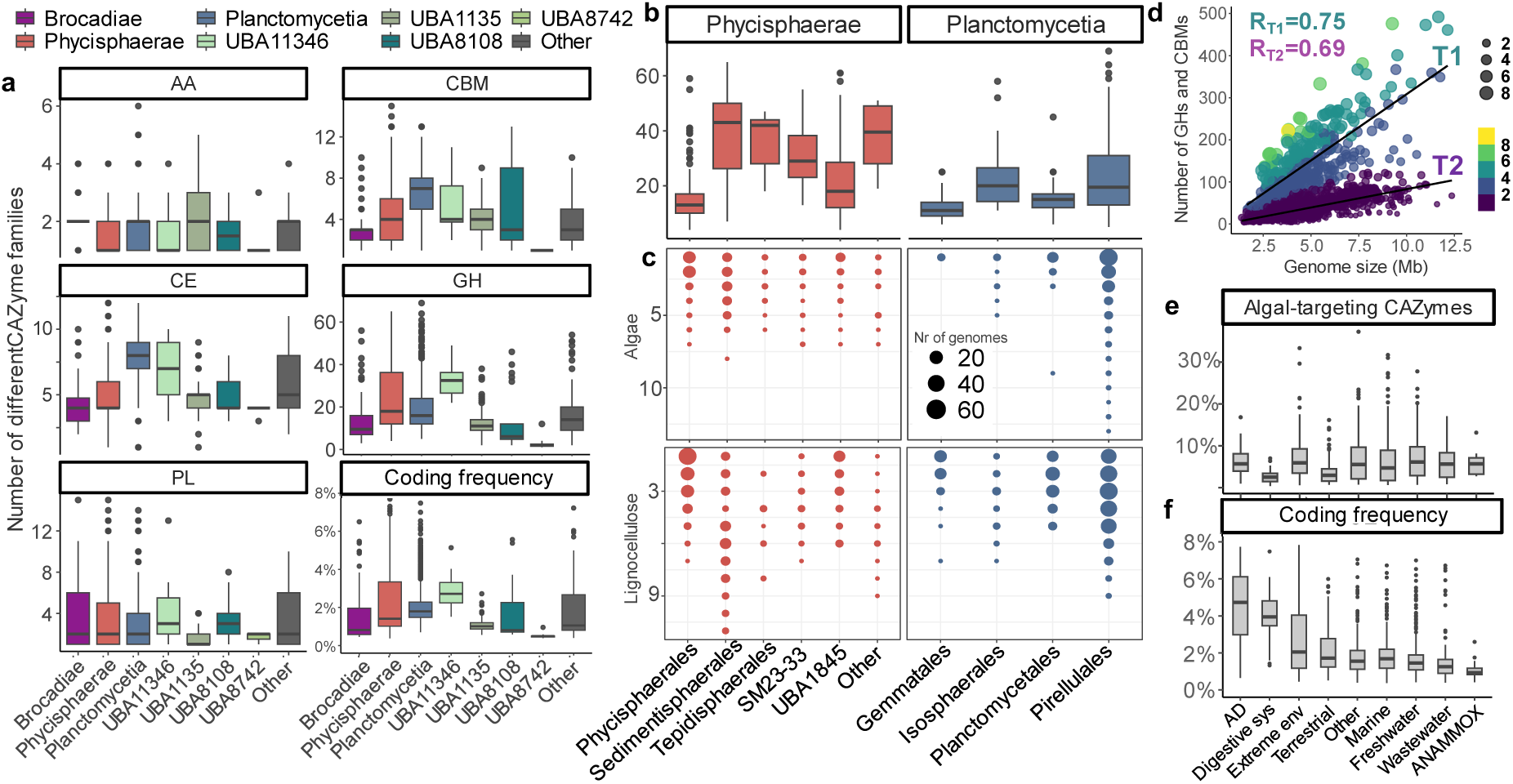
CAZyme family diversity and coding frequency of individual genomes of *Planctomycetota* coloured by class affiliation. Only main trends are shown **a.** CAZyme family diversity at the class level (statistically significant, Kruskal-Wallis test p < 0.05). Last panel: CAZyme coding frequency (percentage of CAZymes versus protein coding genes) at the class level **b.** GH family diversity at the order level for two main classes of *Planctomycetota* **c.** Dotplots presenting number of different GH and PL families with assigned putative functions in each planctomycetotal order **d.** Genome size versus number of GH and CBM coding genes for each planctomycetotal genome, coloured by CAZyme coding frequency. Trends were established based on the thresholds for low (T2, <2%) and medium to high (T1, >2%) coding frequencies. Size of dots and corresponding colours represents the range of CAZyme coding frequency between 0 and 8% **e.** Ratio of algalytic CAZymes encoded by planctomycetotal genomes, grouped at the environmental origin (“habitat”). **f.** CAZyme coding frequency calculated for genomes grouped at the environmental origin (“habitat”).

### Diversity of encoded GHs points to adaptability of individual *Planctomycetota* towards polysaccharides degradation

A broad range of synergistically acting CAZymes is required for the efficient degradation of complex polysaccharides by individual bacteria [49]. Therefore, we examined the potential hydrolytic capacity of individual microbes, by looking at the diversity profiles of CAZymes in genomes assigned to the same phylogenetic class (Fig. 3a) and order (Fig. 3b). The highest diversity of GHs is attributed to the UBA11346 putative class (38 ± 8 GHs per genome) while the lowest GH-encoding potential (between 4 to 13 different GHs) is typical to members of *Brocadiae* and the yet-unclassified classes UBA8108, UBA8742 (Fig. 3a; Additional File 2, Table S7).

Comparing members at the order level, genomes assigned to *Sedimentisphaerales* (*Phycisphaerae* class) display the highest diversity of hydrolysing enzymes assigned to GH and PL families, encoding on average 47 ± 19 and 7 ± 4 distinct subfamilies per genome, respectively (Fig. 3b; Additional File 2, Table S7). In addition to *Isosphaerales* (*Planctomycetia* class; GH diversity 32 ± 16) which are the most recognised planctomycetotal degraders, *Tepidisphaerales* (*Phycisphaerae* class; GH diversity 41 ± 11) commonly found in terrestrial habitats also show multiple distinct GH modules that indicate their capacity to degrade diverse carbohydrates (Fig. 3b). The genomes assigned to *Pirellulales* and *Gemmatales* orders follow with the subfamily diversities of 27 (± 15) and 15 (± 6) respectively.

Accordingly, the *Sedimentisphaerales* order encode the highest number of distinct GH/PL families putatively targeting lignocellulose including xylans (up to 11 families, Fig. 3c bottom panel; Additional File 2, Table S3). Genomes assigned to the SG8-4 putative family are characterised with one of the highest GH/PL diversities among all the *Planctomycetota* (Additional File 7, Fig. S2). Nevertheless, most of planctomycetotal genomes is characterised with at least five separate CAZymes targeting lignocellulose (Fig. 3c). Among other polysaccharide targets, *Pirellulales* encode the highest number of GH/PL targeting algal carbohydrates i.e. up to 13 families (Fig. 3c, top panel).

### Diverse accessory modules support the debranching of complex carbohydrates

Compared to other CAZyme classes, AAs are only occasionally encoded in planctomycetotal genomes (Fig. 3a). Still, certain representatives of the *Planctomycetia* class, encode up to six different AA domains, including putative lignin peroxidases from the AA2 family (Additional File 2, Table S3). They are mainly encoded in unclassified *Planctomycetota* from UBA8742 and UBA1135, equipping these members with plausible capacity to degrade lignin and lignin derivates. All the identified lytic polysaccharide monooxygenases (LPMOs) were assigned to the family AA10, which is the only LPMO family present in bacteria. They were mainly detected in genomes of uncultured members of UBA8108 putative class, the phycisphaeral SM1A02 putative family (present in 25% of genomes) and sporadically in other members of the phylum (see further discussion in Additional File 6).

The representatives of the *Planctomycetia* class display a higher diversity of CE families than other *Planctomycetota* (Fig. 3a), encoding on average 7 to 9 (± 2) different CE families per genome. However, certain members of marine *Phycisphaerales* encode even up to 12 CE families, which is comparable to genomes affiliated to *Planctomycetales* and *Pirellulales,* both representing the *Planctomycetia* class (Additional File 2, Table S7). Overall, *Planctomycetales* and *Pirellulales* trend towards a higher number of esterases including CEs and sulphatases, critical enzymes for debranching algal polysaccharides (Additional File 7, Fig. S3).

The diversity of GH families well reflects the variety of CBM modules in planctomycetotal genomes (Additional File 2, Table S8). Accordingly, *Isosphaerales* (rho = 0.70) as well as phycisphaeral *Sedimentisphaerales* (rho = 0.82), UBA1845 (rho = 0.76) and SM23-33 (rho = 0.70) show a strong correlation (p < 0.05) between GH and CBM family diversity. Interestingly enough, members of *Tepidisphaerales* having one of the highest GH diversities do not follow this trend (rho = 0.21, p = 0.49). High corelation values could also result from the fact that some GH enzymes contain additional domains accommodating other CAZyme modules, especially CBMs, forming multi-modular enzymes [5]. For instance, members of *Sedimentisphaerales* encode on average 10% of CAZymes with multi-modular characteristics (Additional File 7, Fig. S4).

At present, planctomycetotal CAZymes remain largely unexplored. Therefore, we estimated the novelty of planctomycetotal CAZymes by comparing their sequences to the entries in the CAZy database [4] as well as we evaluated the abundance of unclassified CEs, GHs, and PLs that could potentially constitute novel functionalities. Overall, CAZymes encoded in *Planctomycetota* are moderately related to other bacterial CAZymes with the protein sequence identity oscillating on average between 48% and 62%, and some groups such as *Gemmatales* or UBA8108 putative class encode the highest ratio of unclassified CAZymes (further discussed in Additional File 6).

### CAZyme gene coding frequency is governed by phylogeny and is genome size-dependent

In general, genome size correlates positively with the number of genes engaged in carbohydrate metabolism [50]. Different *Planctomycetota* orders are characterised by significantly different CAZyme gene coding frequencies i.e. a percentage of CAZymes compared to the other protein-coding genes (Fig. 3, “Coding frequency“). nterestingly, although an average phycisphaeral genome is smaller than genome representing *Planctomycetia* class (Fig. 1c), members of both classes have similar CAZyme coding frequencies (Fig. 3a). Looking at individual genomes, the highest CAZyme coding frequency was attributed to an uncultivated member of the *Thermoguttaceae* family within the *Pirellulales* order (9.1%), followed by the SG8-4 putative family genome (8.7%) and an *Anaerohalophaeraceae* member (8.3%), with the latter two representing *Sedimentisphaerales* (Additional File 2, Table S9). These genomes were retrieved respectively from the ruminant gastrointestinal tract system (*Thermoguttaceae*) and lab-scale anaerobic digestion studies (*Sedimentisphaerales*), indicating the importance of high CAZyme gene content to the microbial metabolism in carbohydrate-rich environments. Interestingly, genomes assigned to the putative order of *Brocadiae* class (but not *Brocadiales*) were also characterised with higher CAZyme coding frequency (6.7%) as well as some members of unclassified *Planctomycetota* (up to 7.7%, “Other” class).

Coding frequency of planctomycetotal AAs, CEs and PLs clearly correlates with the genome size of bacteria (R = 0.73, p < 0.01, Additional File 7, Fig. S5) but the same could not be observed for GHs and CBMs (R = 0.5, p < 0.01). Instead, GH+CBM coding frequency is shown to follow two main trends (T1 and T2), further illustrated in Fig. 3d. For the same genome size, T2 bacteria are characterised with lower GHs and CBMs coding frequency than T1. The majority of T2 (n = 1053) represents *Planctomycetota* belonging to the *Brocadiae*, putatively novel classes and *Planctomycetia* (except *Pirellulales*) and partially *Phycisphaerae* (vastly including *Phycisphaerales*) classes. Highly packed with CAZymes, T1 *Planctomycetota* are much less numerous (n = 160), and they are primarily assigned to the *Sedimentisphaerales* and *Tepidisphaerales* orders of *Phycisphaerae* and UBA11346 class as well as certain families of the *Pirellulales* order, mainly *Lacipirellulaceae* and *Thermoguttaceae* (Additional File 2, Table S9). Interestingly, T1 genomes are also characterised with more similar CAZyomes in between each other as revealed in the clustering analysis using Dirichlet multinomial mixture models (Additional File 2, Table S9). *Planctomycetota* following T1 were majorly sourced from environments where anaerobic conditions prevail, including extreme habitats, engineered and animal digestive systems where indeed the highest CAZyme coding frequencies are observed (Fig. 3f).

### Functional coupling of CAZymes could help coordinate complex polysaccharide degradation in different *Planctomycetota*

Certain bacteria tend to cluster their CAZymes with complementary functions into so-called CAZyme gene clusters (CGCs). Arrangement in clusters allows to coordinate gene expression leading to the protein ensembles required for a complex carbohydrate saccharification [46]. The best studied example is the polysaccharide utilisation locus (PUL) of *Bacteroidota* [51], however, similar gene clusters were also discovered in other bacterial phyla [52]. To the best of our knowledge, CAZyme clusters have not been functionally characterised in *Planctomycetota* so far. Our genomic analysis revealed that the majority of members belonging to *Brocadiae*, *Phycisphaerae* and putative UBA11346 classes co-localise on average more than 50% of their GHs (Additional File 2, Table S10; Additional File 7, Fig. S6). For comparison, up to 51% of predicted GHs are clustered in *Bacteroidetes cellulolysiticus*, which represents one of the highest scores in bacterial domain [53]. Knowing that some planctomycetotal genomes in our database are incomplete and fragmented (Fig. 1b), the number of CAZymes within gene clusters is certainly higher.

The CAZymes targeting different fractions of lignocellulose are regularly found within CGCs in almost all planctomycetotal classes, except some unclassified *Planctomycetota* i.e. UBA11346 and “Other” class members, whose genomes significantly co-localise only glucan-targeting CAZymes (Fig. 4a). Pectinolytic CAZymes are significantly more co-localised only in members of *Planctomycetia* and *Phycisphaerae* classes, while in contrast, all the glucan-targeting CAZymes are clearly found within CGCs in all *Planctomycetota* (Fig. 4a). Recently, in genomes with few GHs, most of the GH coding genes were shown to be scattered [46]. This is in contract to our observations, as *Brocadiae* genomes are characterised with the lowest CAZyme coding frequency and diversity among studied *Planctomycetota*, at the same time they cluster most of their GHs assigned to α-glucanases (Fig. 2e; Fig. 4a; Additional File 4, Table S2).

**Fig. 4.**
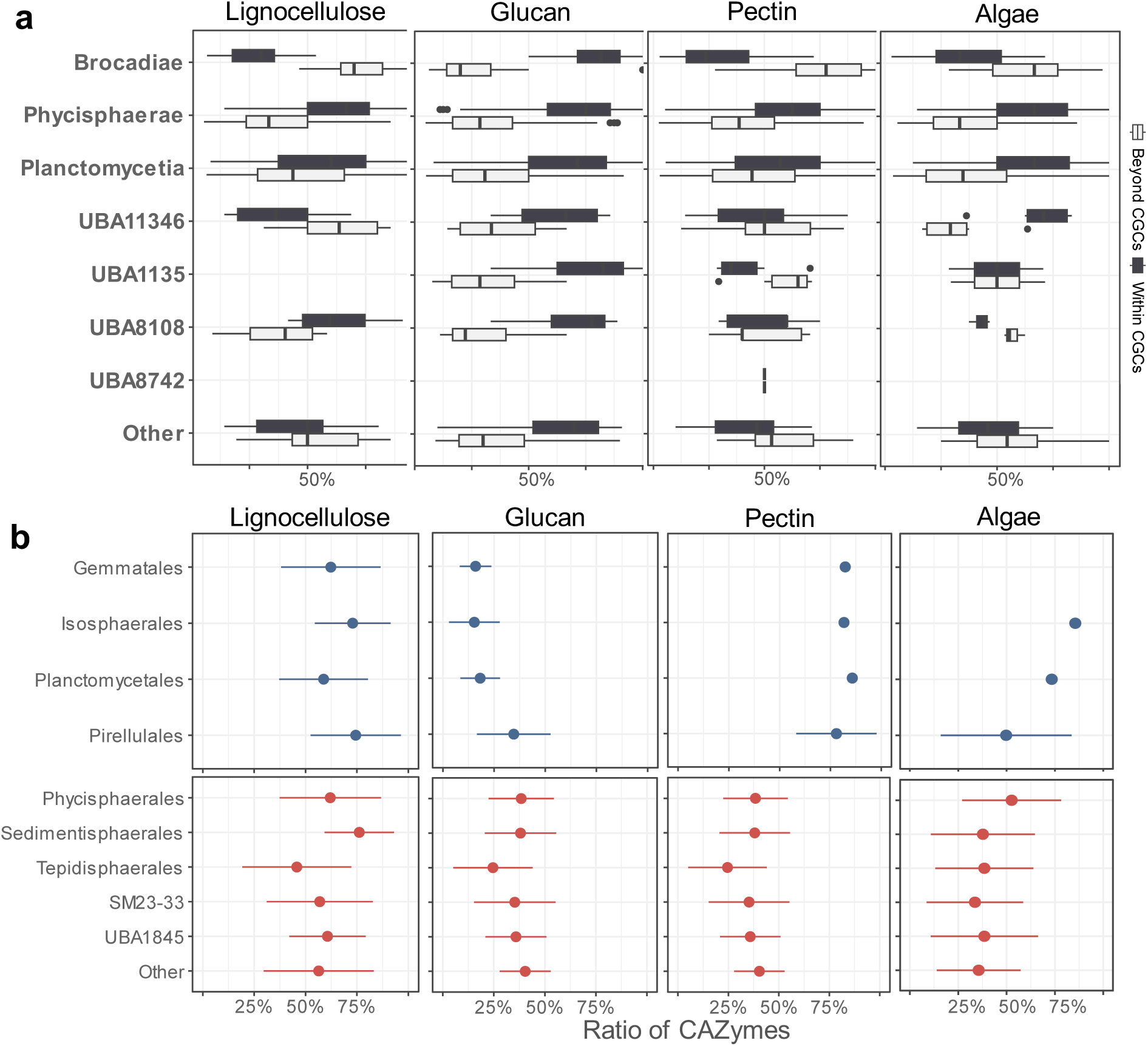
CAZyme gene clustering and predicted localisation of planctomycetotal CAZymes putatively engaged in the degradation of specific polysaccharides **a.** GH and PL families within or beyond CGCs involved in the degradation of lignocellulose (cellulosic and hemicellulosic fractions), glucans (α-glucans), pectins and algal-derived polysaccharides, estimated for class (b) and order (c) levels. In light grey CAZymes outside CGCs. **b.** Ratio of lignocellulolytic, glucanolytic, pectinolytic and algal-targeting CAZymes with any type of predicted signal peptide of the *Planctomycetia* and *Phycisphaerae* orders.

In continuation, we looked at the ratio of CAZymes within CGCs in members of orders from *Planctomycetia* and *Phycisphaerae* classes. While genomes of *Isosphaerales* and *Pirellulales* frequently co-localise their CAZymes, *Gemmatales* and *Planctomycetales* (*Planctomycetia* class) do not encode significantly more co-localised lignocellulolytic, pectinolytic and algalytic CAZymes (p < 0.01) (Additional File 7, Fig. S7). The CAZyme families frequently encoded within CGCs for members of the *Planctomycetia* class are predominantly GH13, GH33 and GH165 and a wide array of enzymes targeting ester linkages, such as CE1, CE14, CE15 (Additional File 7, Fig. S8). Among phycisphaeral orders, most of their CAZymes targeting glucans, pectins, lignocellulosic and algal polysaccharides are co-localised (p < 0.01), including common GH2, GH10, GH29, GH57, and GH78 (Additional File 7, Fig. S7). A similar trend to unclassified *Planctomycetota* UBA11346 can be observed in genomes of SM23-33 putative order. Members belonging to this group preferentially encode CAZymes scattered throughout their genomes, except the α-glucanases which are shown to be significantly more co-localised (Additional File 7, Fig. S9). The genomes of SG8-4 putative family placed within *Sedimentisphaerales* co-localise most of their CAZymes (Additional File 7, Fig. S10), including a high fraction of CGCs consisting of potentially novel, functionally unknown GHs (Additional File 2, Table S11).

### The presence of signal peptides indicates a large portion of planctomycetotal CAZymes are potentially extracellular or membrane-bound enzymes

Extracellular enzymes play an important role in initiating the hydrolysis of complex carbohydrates to shorter oligosaccharides, which are small enough to be transported across the cell membranes for further cytoplasmic sugar utilisation [54]. Microbial mechanisms to secrete enzymes involve among others Sec and TAT pathways in which proteins are flanked with N-terminal short peptides called signal peptides (SP) [55]. To estimate how many of *Planctomycetota* CAZymes can be putatively secreted, we verified whether their enzymes harbour N-terminal SPI (Sec pathway) or Twin Arginine Transport (TAT) peptide signals. Majority of *Planctomycetota* classes were indeed predicted to harbour Sec or TAT pathway signal peptides in more than 50% of total number of CAZymes (Additional File 7, Fig. S11). Among all *Planctomycetota*, putatively secreted are predominantly CBMs, while the proportions of extracellular CAZymes belonging to CEs, GHs and PLs significantly varies across different planctomycetotal groups (Additional File). Looking at the other types of signal peptides appended to the planctomycetotal CAZymes, we could predict that most of the planctomycetotal classes, encode in their genomes CAZymes with lipoprotein signal peptides cleaved by Lsp/SPase II (leader peptidase or signal peptidase II, Additional File 7, Fig. S11).

Subsequently, we assessed the ratio of planctomycetial and phycisphaeral CAZymes with any type of signal peptides according to the different substrate specificity (Fig. 4b). Lignocellulolytic CAZymes are regularly flanked with signal peptides and accordingly, order representatives belonging to *Sedimentisphaerales, Pirellulales,* and *Isosphaerales* encode on average 76.1% (± 16.9), 74.4% (± 21.9), 72.9% (± 18.4) of these putatively extracellular enzymes, respectively. In turn, CAZymes targeting α-glucans are sporadically predicted to possess any of the signal peptides and members of *Phycisphaerales* encode on average the highest ratio of putatively secreted α-glucanases i.e. 38.4% (± 15.9). The profound difference between classes can be observed in pectinolytic and algalytic CAZymes (Fig. 4b). Members of orders assigned to *Planctomycetia* such as *Planctomycetales* (85.7% ± 19.9), *Gemmatales* (82.2% ± 22.8) and *Isosphaerales* (81.6% ± 20.4) encode the highest ratio of putatively secreted pectinases. Despite low number of algal-targeting CAZymes in the genomes belonging to *Isosphaerales* (85.5% ± 18.8), most of them are predicted to be produced extracellularly, while *Pirellulales* putatively secrete on average a half of the encoded algalytic repertoire (49.7% ± 33.8).

## Discussion

In this study, we investigated the metabolic potential of *Planctomycetota* towards polysaccharides degradation. In the absence of cultivable representatives for many of the phylogenetic groups, bioinformatic analyses allow us to characterise microbial metabolisms *in silico*. Nevertheless, a better understanding of planctomycetotal metabolisms is further required to successfully implement their future biotechnological utilisation in the developing biorefinery sector. We hope that the findings presented in our study draw the attention of the scientific community to the *Planctomycetota* phylum as a new and valuable source of CAZymes in the context of both ecological and biotechnological relevance.

### *Planctomycetota* as a reservoir of biotechnologically relevant enzymes

The complete deconstruction of polysaccharides requires GH interaction with other CAZymes, including PLs responsible for non-hydrolytic cleavage of glycosidic bonds, carbohydrate esters hydrolysing CEs, as well as other redox enzymes with auxiliary activities such as AAs, including LPMOs [56]. Collectively, the planctomycetotal enzymatic machinery is a promising alternative for designing novel tools in the context of the biofuel production processes. Members of this phylum encode nearly 80% of currently known GH families, likely reflecting the wide range of possible targets. Given that high diversity of CAZymes advocate for greater capacity of microorganisms to hydrolyse a wider range of complex polysaccharides in diverse environments [49], members of unclassified UBA11346, planctomycetial (*Isosphaerales, Pirellulales*), and phycisphaeral (*Sedimentisphaerales, Tepidisphaerales*) orders demonstrate the highest ability to target polysaccharides including lignocellulosic and algal-derived polysaccharides. Therefore, untapped CAZyme diversity of *Planctomycetota* is a potent reservoir to identify GH families selective for specific linkages of rare polysaccharides that might not be utilised by other microorganisms, serving as compelling target for defined biotechnological applications.

It has been suggested LPMOs efficiently degrade the recalcitrant lignocellulose fractions of the biomass, boosting the overall activity of common GHs [57, 58]. The simplification of enzymatic cocktails would significantly reduce the cost of the enzymatic biomass processing technologies, e.g. for bioethanol production, therefore representing industrially relevant enzymatic preparations. Interestingly, in the course of our analysis, we identified LPMO coding genes in *Planctomycetota* genomes, mainly representing uncultured members of UBA8108 putative class, the phycisphaeral SM1A02 putative family and sporadically in other members of the phylum. So far, none of the planctomycetotal LPMOs have ever been analysed, therefore their enzymatic activities should be further studied to determine their effectiveness for biomass deconstruction (see further discussion in Additional File 8).

Multi-modular enzymes often show multi-activity capacities, allowing to simultaneously target diverse types of carbohydrates, thus representing biotechnologically relevant targets [52, 59]. Indeed, beside a clear correlation between number of GHs and CBMs for some of the analysed planctomycetotal orders, we could also observe that multiple members of *Planctomycetota*, especially *Sedimentisphaerales*, encode a high ratio of multi-modular CAZymes. GHs with multiple CBMs are typical for bacteria but complex CAZymes with different enzymatic modules are relatively less often found within bacterial genomes [5]. Accordingly, within *Sedimentisphaerales,* genomes belonging to the *Anaerohalosphaeraceae* family encode even up to six different modules of putatively hemicellulolytic enzymes, especially involving GH10 modules. Since the conjugation of GHs and CBMs assures higher enzyme affinity to targeting substrates, CAZyme enzymatic complexes might indicate an adaptation strategy e.g., in the competitive environments [60]. Multi-modular CAZymes are also interesting target for the industry market, given that a single enzyme might sustain diverse catalytic modules which likely results in optimal synergy for biomass hydrolysis [61].

### Planctomycetotal polysaccharide utilisation specialists versus generalists

Our analysis indicated that some members of *Planctomycetota* are characterised with less diverse CAZyome, thus could potentially specialise in targeting specific polysaccharide fractions (so called “specialists”). In comparison, genomes with broader CAZyme diversity, could indicate consumers of a wider range of polysaccharides (so called “generalists”). For instance, despite the low GH diversity of *Brocadiales*, their genomes co-localise most of α-glucanases, putatively representing a highly specialised system for the storage polysaccharide metabolism (e.g., glycogen). A limited CAZyme repertoire may also result from the fact that carbohydrates are not the primary source of energy for this largely chemolithotrophic group of bacteria, as they might utilise some other compounds to sustain their metabolism [45, 62]. Members of not yet described families within *Phycisphaerales* (*Phycisphaerae* class) and *Planctomycetales* (*Planctomycetia* class) are also narrow in total CAZyme diversity, however, order members of both classes encode the highest number of CAZyme families assigned with rare functionalities, e.g. enzymes with accessory functions. These findings highlight members of *Phycisphaerales* and *Planctomycetales* orders as interesting candidates for specialised biotechnological applications, where defined utilisation of specific carbohydrate types is of interest [63, 64].

Overall, the CAZyme diversity of *Isosphaerales, Sedimentisphaerales* and *Tepidisphaerales* was vastly similar to members of the *Bacteroidota* phylum, which are typically recognised as generalist carbohydrate degraders [10, 65]. In line with this observation, culturable isolates of both *Isosphaerales* and *Tepidisphaerales* were previously recognised as rich source of diverse hydrolytic enzymes [20], likewise the genomes in our analysis placed within these taxonomic groups. Strains of *Isosphaerales* isolated from boreal peat bogs and *Sphagnum*-dominated wetlands were shown to metabolise a wide variety of complex carbohydrates, including xylans, celluloses, lichenans or pectins [13]. Surprisingly, isolates of *Planctomycetota* from hypersaline habitats representing *Sedimentisphaerales* order (*Phycisphaerae* class) have not so far been shown to utilise complex carbohydrates, presumed to be involved in the degradation of partially debranched glycans [69]. Regardless, genomes in our database placed within this order have overall broad CAZyme diversity, pointing to the high heterogeneity of carbohydrate catabolism between the members of this largely uncharacterised planctomycetotal group. Of particular note is the SG8-4 putative family of *Sedimentisphaerales* found in diverse environments including aquatic and engineered (e.g. AD) systems [69], whose members display one of the highest genomic CAZyme diversity and functional versatility among all *Planctomycetota*. Nevertheless, current exploitation of this planctomycetotal group is limited as no cultured representatives are yet available.

### Planctomycetotal CAZyme repertoire is shared by the members of the phylogenetic groups and unique environmental conditions

Our analysis signifies the importance of phylogeny rather than habitat as a main driver for CAZyome composition, consolidating the previous observations for other bacterial phyla [10]. Overall, *Planctomycetota* from diverse environments encode similar average potential towards different types of biomasses, but preferentially encoding CAZyme families specific to certain fractions of polysaccharides. As such, *Phycisphaerales* trend towards higher ratio of enzymatic machinery involving NAG- and algal-targeting CAZymes. This observation is reasonable, considering that almost 70% of all the *Phycisphaerales* genomes in our database originate from aquatic environments where crustacean (chitin) and algal biomass is prevalent [70]. In turn, *Pirellulales, Sedimentisphaerales* and *Tepidisphaerales* encode a considerable enzymatic machinery dedicated to breaking down pectins, which are a significant component of the plant cell wall matrix [71]. They also show an increased potential for oligosaccharide deconstruction what characterises the degraders that target multiple and diverse polysaccharides [49]. *Pirellulales* genomes assigned to the *Pirellulaceae* family, are characterised with a much higher GH family diversity than the order average, constituting interesting “outliers” possibly targeting wider range of polysaccharides. Considering a rich selection of algal-degrading enzymes of *Pirellulales* as well as high number of CEs and sulphatases encoded in their genomes, we could infer the presence of a system designed to scavenge the algal biomass. This reinforces the earlier observations of different *Pirellulaceae* members, predominantly assigned to *Rhodopirellula*, to live in close association with macroalgae and assumed to possess a wide repertoire of CAZymes attacking sulphated carbohydrates [72, 73].

A higher number of functionalities encoded in bacterial genomes was proposed to make up the larger genome size, shifting the potential for the discovery of new functionalities towards bigger genome species [50]. Previously, a positive correlation between the planctomycetotal genome size and the number of biosynthetic gene clusters (BGCs) was observed [74]. Here, we observed an unexpected tendency of *Planctomycetota* to discriminate into two main trends owing to the different number of encoded GHs and CBMs among the bacteria within the same range of genome size (which we tentatively called T1 and T2 *Planctomycetota*). As we further realised, genomes characterised with the higher trend (T1) were commonly retrieved from extreme habitats, animal gut and anaerobic digesters – all representing environments with conserved and stable conditions [75, 76]. In general, bacteria found in multiple environments shift towards larger genomes, as opposed to members of specific niches, characterised with reduced genomes due to loss of unnecessary genes [77]. Such specialisation has already been observed in certain lineages of *Planctomycetota* transitioning from sediment/soil to freshwater environments [78]. Nevertheless, genome streamlining is a common route for microorganisms inhabiting oligotrophic environments but with limited significance in nutrient-rich habitats [79]. The genomic expansion could involve the acquisition of additional genes coding for carbohydrate metabolism that possibly allow T1 *Planctomycetota* to capitalise on the resources present in the dwelled environments. Furthermore, genomes following T1 trend also share their CAZyme pool, and have a tendency to higher CAZyme coding frequency and diversity compared to other *Planctomycetota* (T2 trend). Ultimately, this is a known observation for microorganisms from AD, animal digestive tract and extreme environments, which were previously recognised as promising spots for the discovery of biomass-degrading enzymes suitable for biofuel production [8, 80, 81].

### Different potential strategies of *Planctomycetota* for complex polysaccharide deconstruction

Distinct groups within *Planctomycetota* phylum encode co-localised CAZymes like other well-studied bacteria such as *Bacteroidota,* widely recognised for their PUL diversity [82–84]. Throughout our analysis, genomes assigned to *Sedimentisphaerales* recur persistently in the different contexts of carbohydrate utilisation. This order encompasses the SG8-4 putative family which genomes seem to co-localise most of their CAZymes into operons likely involved in complete degradation of polysaccharides. Closer examination of genomes belonging to this order show high encoded degradative potential towards sulphated N-glycans and different hemicelluloses. Members of planctomycetotal orders may employ potentially distinct strategies towards different polysaccharides, which is crucial factor for understanding their ecology, but also a critical aspect in designing future bioprocessing. Among unclassified *Planctomycetota*, members assigned to UBA11346 deserve some attention. Despite low number of genomes in our database (16), they show a wide diversity of encoded CAZyme families and the highest CAZyme coding frequencies. At the same time, their CAZymes are regularly found beyond the CAZyme clusters, potentially representing a different enzymatic strategy than other generalist *Planctomycetota* such as *Isosphaerales, Pirellulales* or *Sedimentisphaerales*.

Most of the planctomycetotal genomes flank their CAZymes with signal peptides which indicates the export of the protein through the membranes. This suggests that bacteria may secrete CAZymes for targeted extracellular degradation of carbohydrates, in particular lignocellulose, pectins and algal-derived polysaccharides. From an ecological perspective, enzyme-secreting bacteria are beneficial to the whole community, as they pre-degrade larger fibres into smaller components, which can be used at the same time by other microbes [54]. Such bacteria are recognised as important common goods producers for the microbial community as opposed to “scavengers” who habitually rely on their products [85].

Nevertheless, it was suggested that certain *Planctomycetota* selfishly import marine polysaccharides through an unknown mechanism [85]. In such case, CAZymes flanked with the N-terminal peptide would only be transported to the periplasm, where the main saccharification could take place. In addition, members of unclassified *Planctomycetota* along with *Phycisphaerae* are predicted to possess lipoprotein signal peptides (Lsp/SPII) appended to their CAZymes. This type of signal peptide often serves for intracellular localisation [37], thus putatively support anchoring their enzymes to either the inner or the outer cell membrane. In comparison with the truly extracellular enzymes, cell-membrane-anchored extracellular CAZymes would benefit more the host than the other community members, allowing for the higher share of the liberated oligosaccharides to be taken by the main enzyme producer [85, 86]. Similar mechanisms were previously described i.e. for a starch utilisation system (sus) of *Bacteroides thetaiotaomicron* [86] and further investigation of comparable systems in *Planctomycetota* is of high priority. An interesting feature, exclusive to *Planctomycetota* was further observed by Boedecker et al. [11], who described an extreme enlargement of the periplasmic space in *Planctopirus limnophila* (*Planctomycetales* order), accompanied by its ability to bind sugar moieties using crateriform structures, when feeding on complex, branched glucan (dextran). Likewise, type IV pili of *Fimbriiglobus ruber* (*Gemmatales* order) were shown to enhance the bacterial adhesion to chitin and other biopolymers [87]. So far, these mechanisms have only been proven experimentally for some species from the *Planctomycetia* class, while it is not known if the same can occur in other *Planctomycetota*. Any further ecological relevance, directly or indirectly engaged in polysaccharide uptake and degradation remains to be scrutinised.

### Limitations of predicting the complex carbohydrate degrading potential through *in silico* approaches

The encoded carbohydrolytic potential of *Planctomycetota* cannot fully reflect the microbial capacity in the environment, and contextualising the genomic measurements to metabolic traits remains difficult [88], hampering our understanding of their ecology at large. It is worth noting that many isolated strains of *Planctomycetota* exhibit slow growth rates [89] and one needs to explore and optimise the culture or strain maintenance procedures before developing specific applications. Culture-based methods are still required for optimising and scaling up biotechnological processes [90], posing an ongoing challenge, however a few methods have already been established for *Planctomycetota* [91, 92]. Even though existing limitations of bioinformatic analysis mitigate our efforts to precisely predict metabolic potential of some microorganisms, they help spotting interesting microbial groups where future isolation attempts should be reinforced [93]. On the other hand, for the discovery of novel CAZymes, metagenomic approaches escapes the bottleneck of culturing microorganisms [56]. We expect to extend the knowledge on carbohydrolytic potential of *Planctomycetota* once more representatives are available in axenic cultures, with special regards to anaerobic yet uncharacterised groups assigned to *Pirellulales* and *Sedimentisphaerales*.

As a general rule, most identified degraders have multiple genes coding for CAZymes [94], however, it does not always serve as a reliable indicator of bacteria with high carbohydrolytic capacity. Such an example are representatives of *Gemmatales* (*Planctomycetia* class), with generally low GH family diversity compared to all the other *Planctomycetota*. However, our findings contrast with the general perception of this order, as bacteria with a broad hydrolytic potential [13]. Previously, cultured representatives of *Gemmatales* were characterised as versatile degraders of multiple biopolymers like xylan, laminarin, lichenin, chitin, and cellulose [21]. Indeed, a significant part of *Gemmatales* genomes is devoted to coding enzymes that could possibly target NAG-based polysaccharides including chitin. However, at the order level, the genomes do not represent a similar genomic potential for lignocellulosic polysaccharides as other members of *Planctomycetota*. Interestingly, a relatively high number of hypothetical CAZymes which currently cannot be classified to any known CAZyme family is revealed in the genomes of *Gemmatales,* likely explaining this phenomenon (discussed in Additional File 8).

The classification of microorganisms is relatively straightforward for groups with well-studied and established taxonomies [26]. However, dealing with novel or poorly characterise taxa may cause uncertainty and current taxonomic placement is likely to change with the discovery and sequencing of new genomes or species. In addition, the limited availability of genomes representing potentially novel *Planctomycetota*, poses a challenge conducting a thorough analysis at the higher taxonomic levels. Nevertheless, in our study we tried to identify the main trends across *Planctomycetota,* narrowing analysis to the order taxonomic level. We argue that within-phylum genome analysis provides a more nuanced and detailed perspective on carbohydrolytic diversity of *Planctomycetota*, enabling us to investigate common patterns that may not be as evident when comparing to other bacterial phyla.

Curiously enough, despite the low number of genomes within *Isosphaerales* in our database (28), half of them represent axenically isolated strains. Conversely, of all 116 recovered genomes that belong to *Sedimentisphaerales,* only four correspond to axenic isolates. This signifies the specific ecological preferences of the different *Sedimentisphaerales* members, which cannot so far be mimicked by commonly used cultivation practices. Nevertheless, the low number of *Isosphaerales* genomes could be explained by the fact that reconstruction of MAGs from highly diverse environments including soil is still largely constrained by current technologies [95]. The emergent underrepresentation of soil microbial MAGs in public databases alleviates our effort to further exploit this genetic resource.

### Perspective for second- and third-generation biomass valorisation with *Planctomycetota*

Despite the general abundance and ubiquity of plant biomass, lignocellulose is an untapped biomass feedstock resource due to its recalcitrance [96]. In consequence, this so-called second-generation feedstock still lacks economic viability at the industrial scale and new approaches are needed to improve the enzymatic hydrolysis of diverse plant biomasses [97].

At present, there is a viable interest in understanding how microorganisms employ different strategies for complex polysaccharide degradation to reinforce the current solutions or develop new industrial technologies [52, 60, 98]. For example, CAZyme gene clusters allow for the coordinated production of multiple, often synergistically acting enzymes [51, 99]. As such, the enzymatic complexes required for complete polysaccharide digestion could be extracted together, simplifying the enzymatic cocktail design [56]. Another example of nature-inspired strategy for biorefinery-related applications is enzymatic pre-treatment of biomass [97]. Extracellular enzymes, secreted directly to the surrounding environment, simplify the process of extraction and downstream processing, often showing advantageous properties compared to enzymes produced intracellularly [100]. In view of all these aspects, we think that members of phycisphaeral *Sedimentisphaerales* and planctomycetial *Pirellulales* characterised with putatively extracellular, diverse and frequently co-localised CAZymes are among the high-priority targets for extending the strategies to be applied to biomass-based biorefineries.

Compared to lignocellulose, algae, which is considered a third-generation biomass, could potentially offer several advantages [101]. The negligible presence of lignin, short generation times and high lipid content make algal biomass less resistant to degradation, reducing the need for energy-intense pre-treatments [102]. Nevertheless, the remarkable variability of unique algal polysaccharides requires new technological platforms that challenge the conventional biorefinery sector [103]. For instance, brown algae seem to be the major obstacle for conventional biorefineries as they produce various fucoidans, the most recalcitrant algal-derived polysaccharides [104]. Given that, it is crucial to design an individual approach for each type of biomass, and nature-inspired cocktails seem to be a promising alternative for the complete conversion of different biomasses to fermentable sugars [105–107]. Initially, we expected *Planctomycetota* retrieved from marine environments to serve as a reservoir for diverse algalytic CAZymes, regarding abundance of algal biomass in seawater. Contrary to our expectations, genomes of *Planctomycetota* retrieved from different environments including engineered systems encode similar potential towards algal biomass than other marine-originated members of this phylum. Furthermore, AD-sourced genomes are significantly enriched in enzymes specific to sulphated glycans, especially fucose-based polysaccharides. Across the phylum, we could detect GH families 29, 107, 139, 151 and 168, but also other polyspecific families that may also exhibit multiple activities towards fucoidans and fucose-containing oligosaccharides such as GH30, GH95, and GH141. Although some members of *Planctomycetota* phylum are already well-known utilisers of sulphated compounds including carrageenans, chondroitin sulphates and fucoidans [19, 85], little is known about the planctomycetotal enzymatic systems involved in the degradation of algal biomass at large. For instance, the complexity of fucoidans pressure bacteria to possess highly specialised enzymatic systems in order to fully degrade them, as described for *‘Lentimonas’* sp. CC4 [104]. Arguably, *Planctomycetota* might also be a key player in degrading various structurally complex fucoidans, given the widespread distribution of CAZymes targeting backbone of sulphated polysaccharides in their genomes.

Overall, the capacity of anaerobic microbes to degrade algal biomass directly in AD reactors opens up a new perspective for its valorisation in the context of biogas production, and other developing biorefineries. As the field of green biotechnology continues to advance, the importance of planctomycetotal-based application is likely to grow.

## Conclusions

The *Planctomycetota* phylum offers a wealth of diverse, potentially novel CAZymes of industrial interest. Our study provides a new perspective on the planctomycetotal carbohydrolytic potential, highlighting the presence of distinct phylogenetic groups with both general and specialised abilities to break down complex carbohydrates. We identified planctomycetotal families affiliated to the *Sedimentisphaerales* and *Pirellulales* orders that are not yet well characterised as suitable candidates for applications in second generation biomass transformation technologies, due to their diverse CAZymes, including extracellular lignocellulose targeting enzymes. In addition, we showed that some *Planctomycetota* possess LPMOs, which can be further employed to boost the overall activity of GHs in lignocellulose hydrolysis. To our surprise, AD-sourced *Planctomycetota* appeared to be well-equipped for degrading algal-derived polysaccharides, thus representing a perspective for a direct algal biomass transformation to bioenergy in methanogenic reactors. Overall, our findings have implications for directing bioprospecting ventures to enable a more effective discovery of CAZymes in *Planctomycetota*. Although the most interesting planctomycetotal models represent still uncultivated bacteria, their enzymes can already be explored for specific applications thanks to their identification and characterisation through *in silico* studies.

## List of abbreviations

CAZyme(s): Carbohydrate Active Enzyme(s)
GH: Glycoside Hydrolases
CE: Carbohydrate Esterases
PL: Polysaccharide Lyases
AA: Enzymes with auxiliary activities
PCoA: Principal Coordinates Analysis

## Availability of data and materials

All data generated or analysed during this study are included in this published article and its supplementary information files. Accession numbers of public genomes used in this study are listed in the Additional File 1, Table S1. Remaining genomes from previous, own studies are available upon request.

## Funding

This study was supported by the National Research Fund, Luxembourg (AFR Grant, ref. 14583934).

## Supplemental Information

**Additional File 1. Table S1.** The initial genome database and final dataset considered in the study. Related metadata: taxonomic affiliation, GC% content, genome size, number of contigs and predicted proteins. **Table S2.** Metadata categories explained.

**Additional File 2. Table S3.** The final count table used in the study: Genome, Gene ID and CAZyme genes based on an HMMER search. **Table S4.** Percentage of genomes encoding each CAZyme family (within phylum). **Table S5.** Percentage of frequent (>90%) and rare (<5%) CAZymes at the order level (selected the most populated orders). **Table S6.** The list of significantly enriched CAZyme families between different planctomycetotal orders. The p values were fadjusted using the Benjamini & Hochberg correction method. **Table S7.** Mean and standard deviation of distinct CAZyme families and subfamilies in each planctomycetotal order. **Table S8.** The correlation between the GH and CBM module number. **Table S9.** The coding frequency for each genome, cluster assignment based on Dirichlet MM and related metadata. **Table S10.** The CAZyme Gene Clusters: the number of CAZyme coding genes that can be found in CGCs. **Table S11.** The CAZyme Gene Clusters in the SG8-4 putative family. **Table S12.** The mean and standard deviation of predicted GH/PL with signal peptides, calculated for planctomycetotal orders. **Table S13.** The CAZyme percentage identity estimated by DIAMOND. **Table S14.** The matrix contains the pairwise percentage identity between all planctomycetotal LPMO sequences.

**Additional File 3. Table S13.** Substrate specificity assigned to the GH and PL families based on [3] and the CAZy database (http://www.cazy.org/).

**Additional File 4. Table S14.** Raw output file from dbCAN2 (overview).

**Additional File 5. Table S15.** Raw output file from CGC Finder.

**Additional File 6. Table S16.** Raw output file from signalP.

**Additional File 7. Supplementary Figures. Fig. S1.** Heatmap presenting the most commonly encoded CAZyme families within planctomycetotal orders (at least 50% of order representatives). Numbers represent a fraction of genomes encoding each CAZyme family (%) belonging to cleavage enzymes (a) and other (b) class of enzymes. **Fig. S2.** CAZyme diversity of selected classes (CE, GH, PL), calculated for individual genomes of Planctomycetota, coloured by class affiliation and grouped at the family level. **Fig. S3.** Encoded percentage of sulfatases (EC number 3.1.6.-) in individual genomes, grouped at order level and coloured by class affiliation. **Fig. S4.** Percentage of multi-modular CAZymes in planctomycetotal genomes. Boxplots show percentage of multimodular CAZymes for each genome at order level coloured by class affiliation. Gray circles show the diversity of multi-modular CAZyme modules (number of unique CAZyme combinations). **Fig. S5.** Number of AAs, CEs and PLs in planctomycetotal genome versus their genome size (Mb). Spearman correlation calculated. **Fig. S6.** A mean fraction of CAZyme coding genes co-localised within CGCs for each planctomycetotal class. **Fig. S7.** CAZyme gene clustering of planctomycetotal CAZymes putatively engaged in the degradation of specific polysaccharides. Only orders of the Planctomycetia and Phycisphaerae classes are shown. GH and PL families within or beyond CGCs involved in the degradation of lignocellulose (cellulosic and hemicellulosic fractions), glucans (α-glucans), pectins and algal-derived polysaccharides. In light grey CAZymes outside CGCs. **Fig. S8.** Common co-localised CAZymes within CGCs for Planctomycetia class. **Fig. S9.** Common co-localised CAZymes within CGCs for Phycisphaerae class. **Fig. S10.** Predicted localisation of CAZymes, coloured at the class level. Only main trends are shown (a) Ratio of CAZymes with predicted SP or TAT signal peptide for each planctomycetotal genome, grouped at the class level (b) Ratio of CAZymes with predicted signal peptides for lipoproteins for each planctomycetotal genome, grouped at the class level.

**Additional File 8. Supplementary Discussion. Fig. S11.** The novelty and uniqueness of planctomycetotal CAZymes. Percentage of CAZymes within planctomycetotal genomes not assigned by DIAMOND to any known CAZyme family, grouped at class (a) and order (b) taxonomic level. **Fig. S12.** Phylogeny of 57 planctomycetotal AA10 protein sequences (coloured in blue and brown) and 15 other AA10 enzymes with confirmed lytic oxidoreductase activity (coloured in black). Neighbor-Joining tree was built using aligned sequences by muscle algorithm. In brackets percentage identity of each protein and its closest blast hit (NCBI entry). Nodes coloured in brown represent proteins with the highest similarity to other *Planctomycetota* (*Poriferisphaera corsica* KS4). Nodes coloured in blue point to other donor bacteria such as *Actinomycetota, Bacillidota* and *Proteobacteroidota.* Nodes coloured in black represent enzymes already described in the literature. **Fig. S13.** The phylogeny of planctomycetotal AA10 protein sequences retrieved from genomes assigned to SM1A02 putative family and the additional 15 AA10 protein sequences from bacteria representing other phyla with the lytic polysaccharide monooxygenase activity described (LPMO). Bootstrap values are shown on branches.

## Notes

### Competing Interest Statement

The authors have declared no competing interest.

