## Additional File 7 for "Comparative genomic analysis of *Planctomycetota* potential towards complex polysaccharide degradation identifies phylogenetically distinct groups of biotechnologically relevant microbes"

**Supplementary Figures**

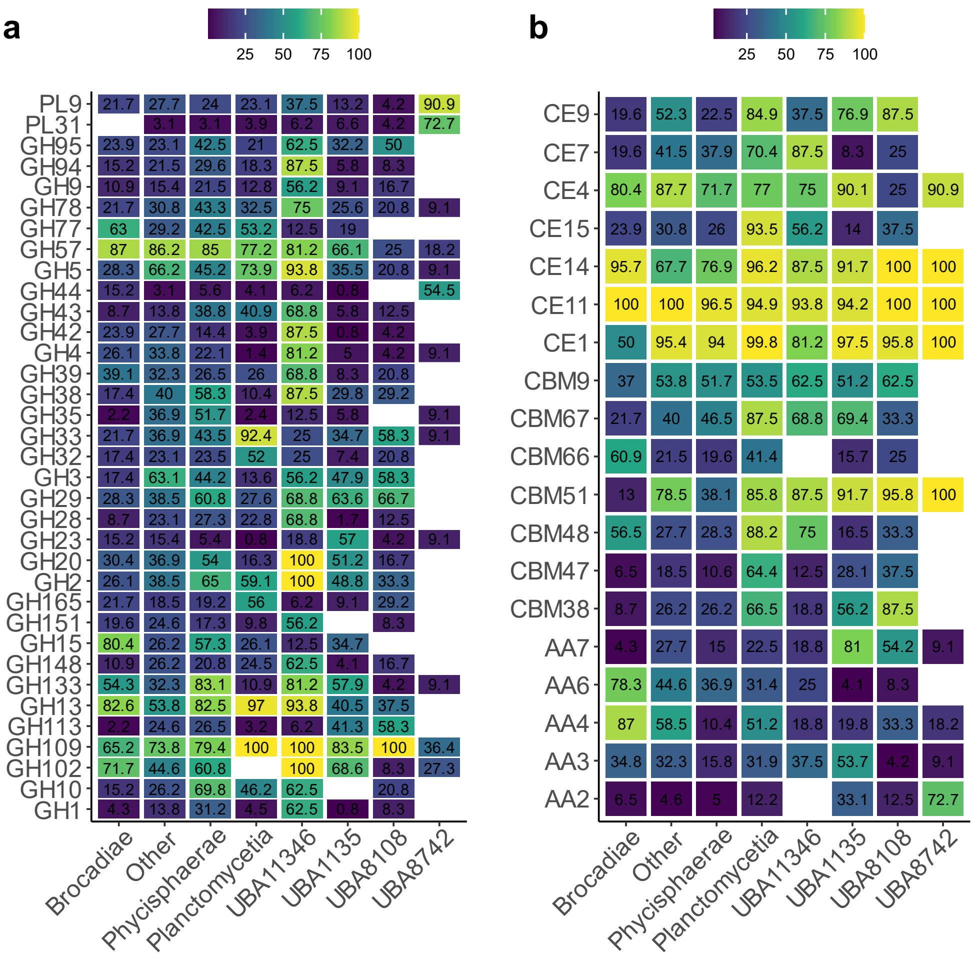


**Fig. S1.** Heatmap presenting the most commonly encoded CAZyme families within planctomycetotal orders (at least 50% of order representatives). Numbers represent a fraction of genomes encoding each CAZyme family (%) belonging to cleavage enzymes (a) and other (b) class of enzymes.

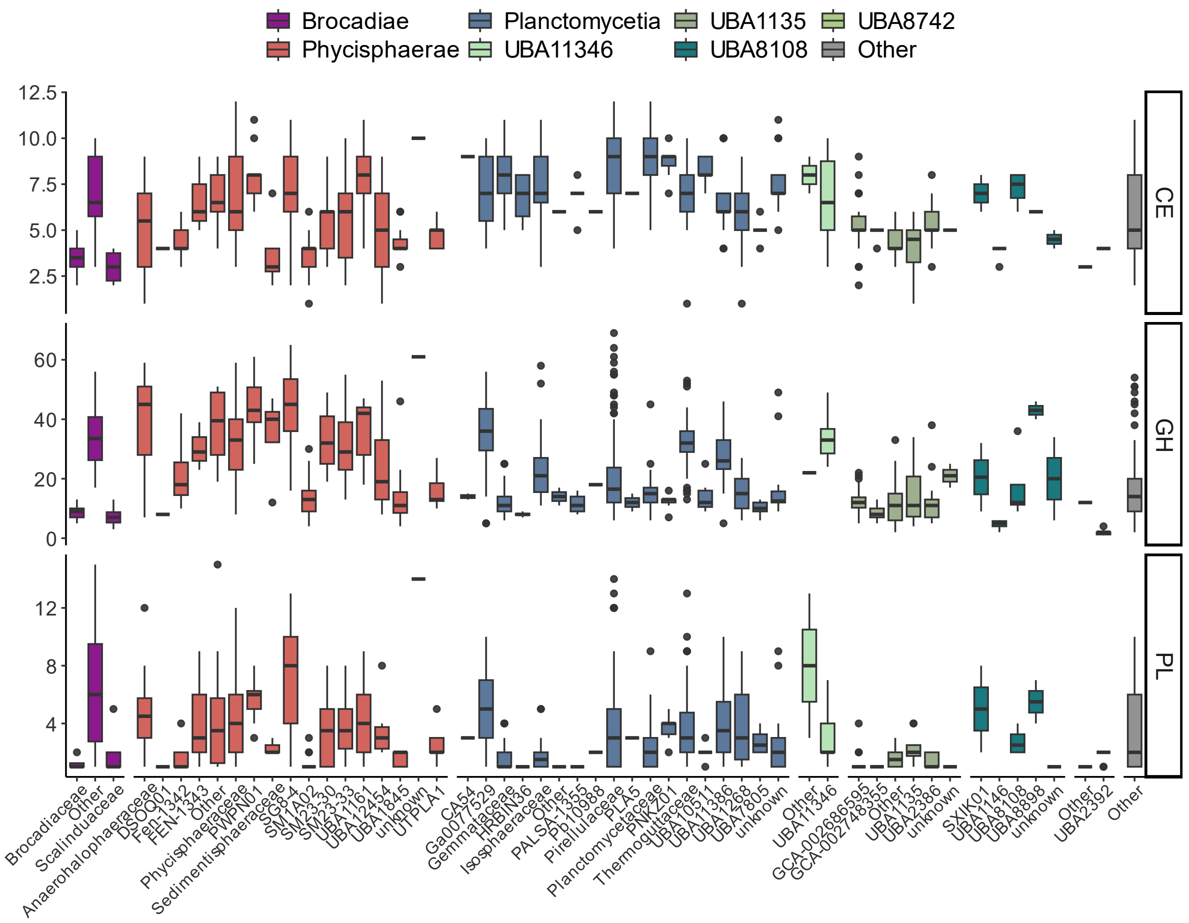


**Fig. S2.** CAZyme diversity of selected classes (CE, GH, PL), calculated for individual genomes of Planctomycetota, coloured by class affiliation and grouped at the family level.

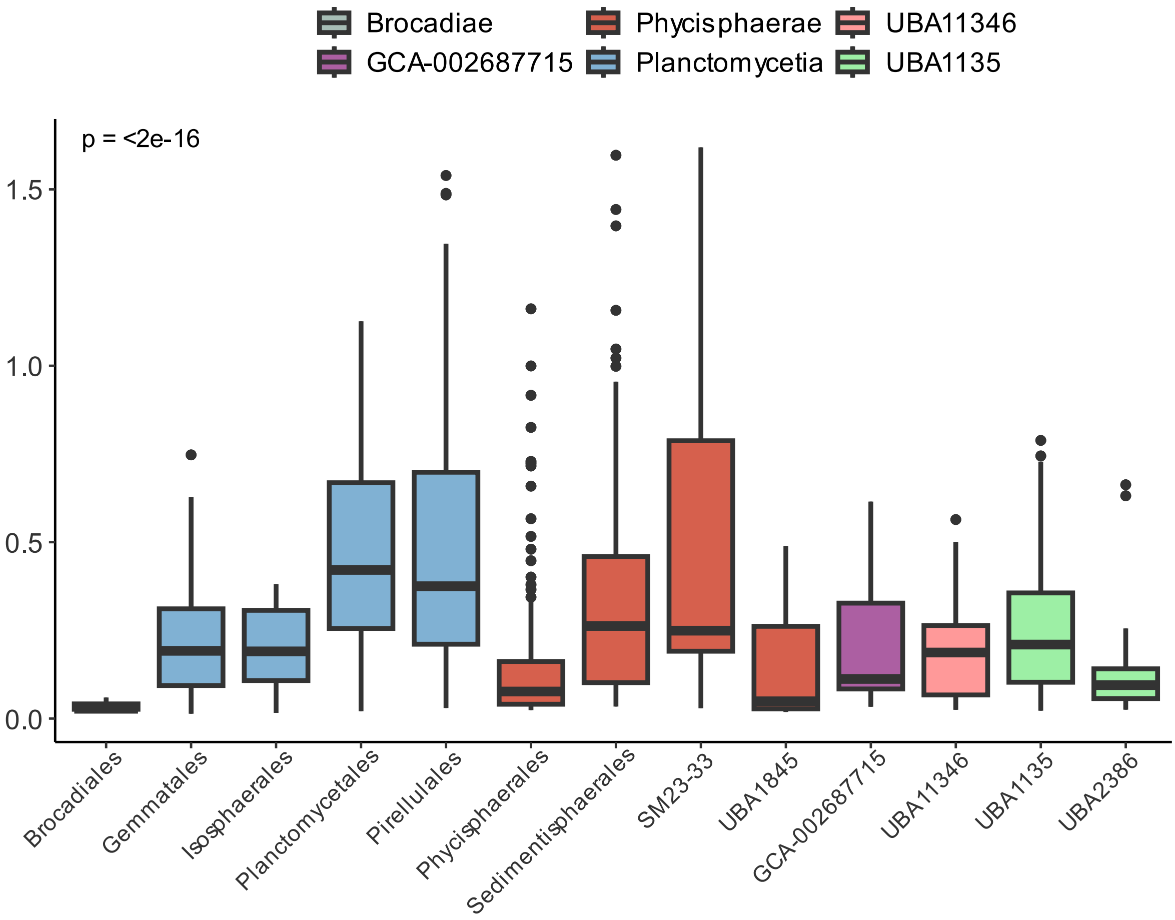


**Fig. S3.** Encoded percentage of sulfatases (EC number 3.1.6.-) in individual genomes, grouped at order level and coloured by class affiliation.

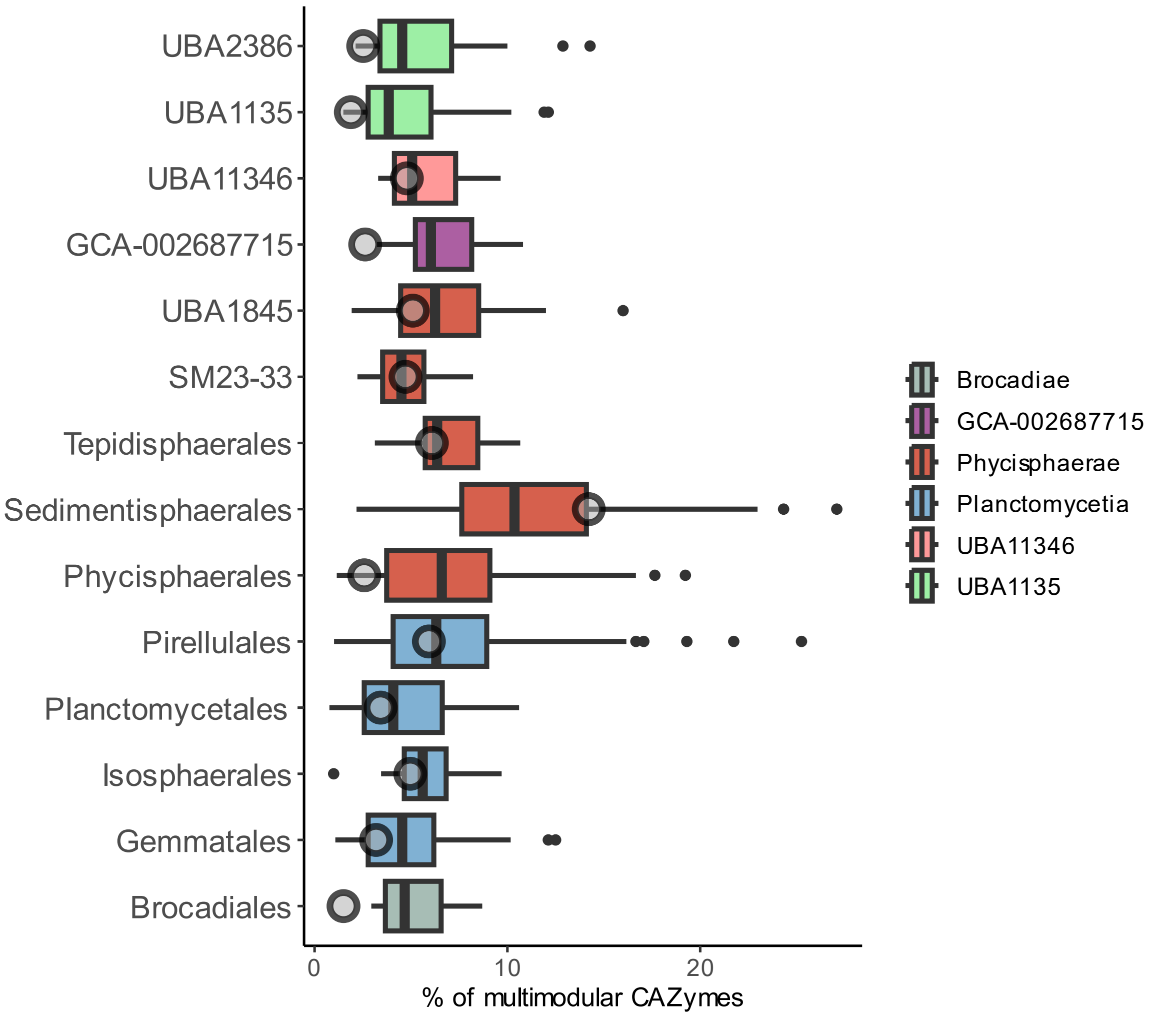


**Fig. S4.** Percentage of multi-modular CAZymes in planctomycetotal genomes. Boxplots show percentage of multimodular CAZymes for each genome at order level coloured by class affiliation. Gray circles show the diversity of multi-modular CAZyme modules (number of unique CAZyme combinations).

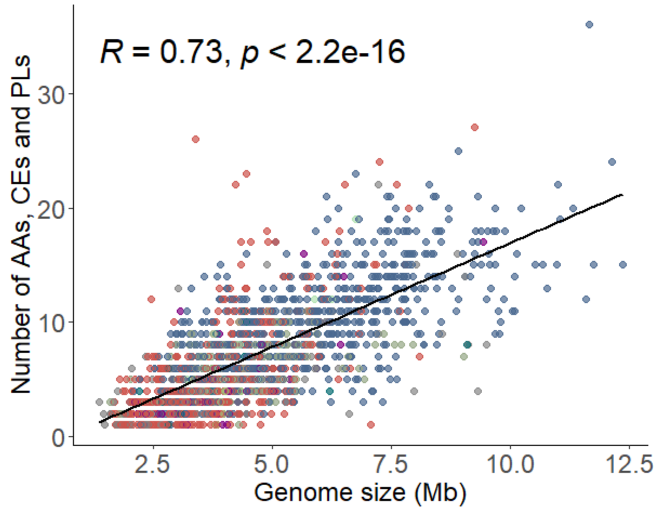


**Fig. S5.** Number of AAs, CEs and PLs in planctomycetotal genome versus their genome size (Mb). Spearman correlation calculated.

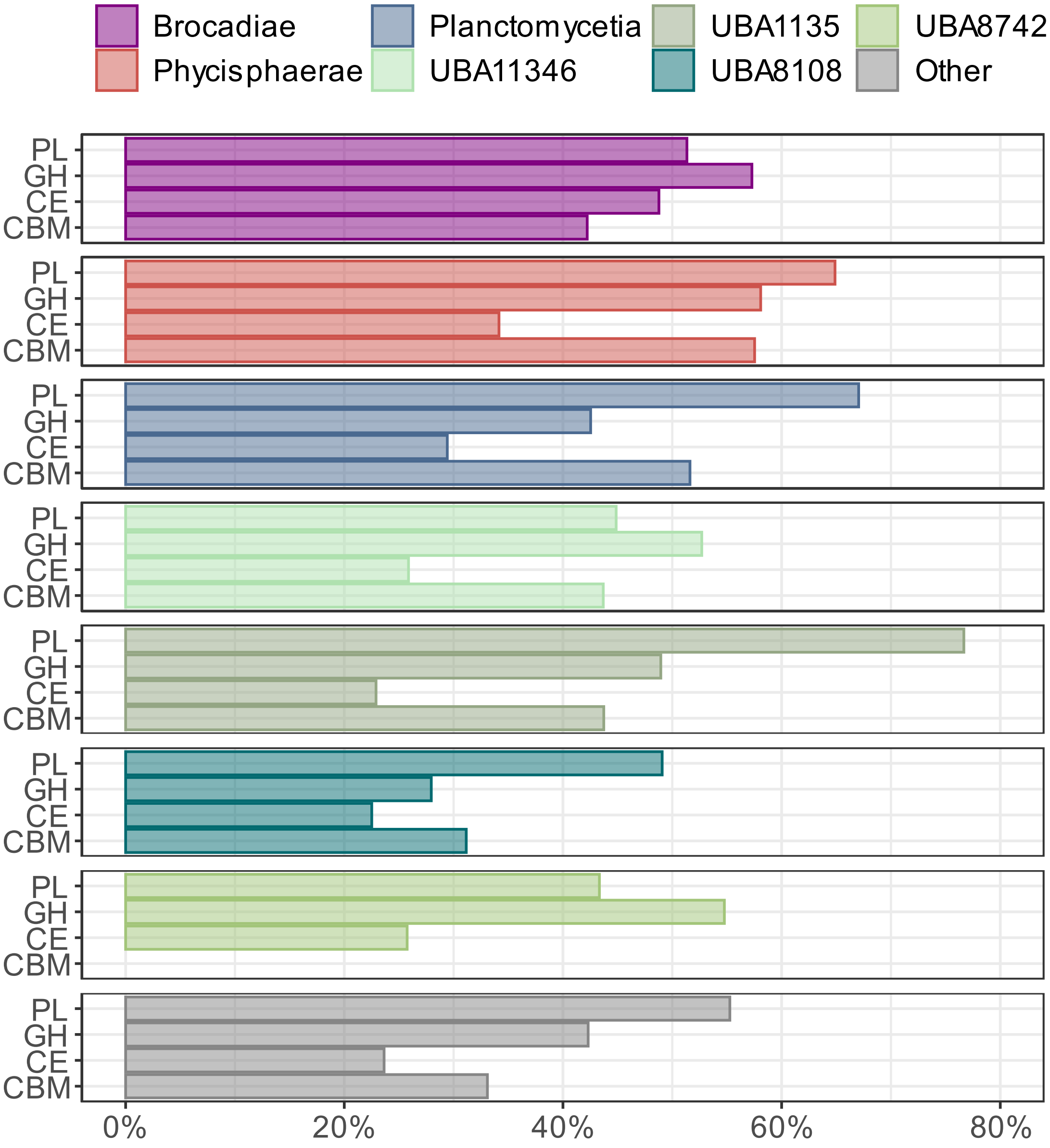


**Fig. S6.** A mean fraction of CAZyme coding genes co-localised within CGCs for each planctomycetotal class.

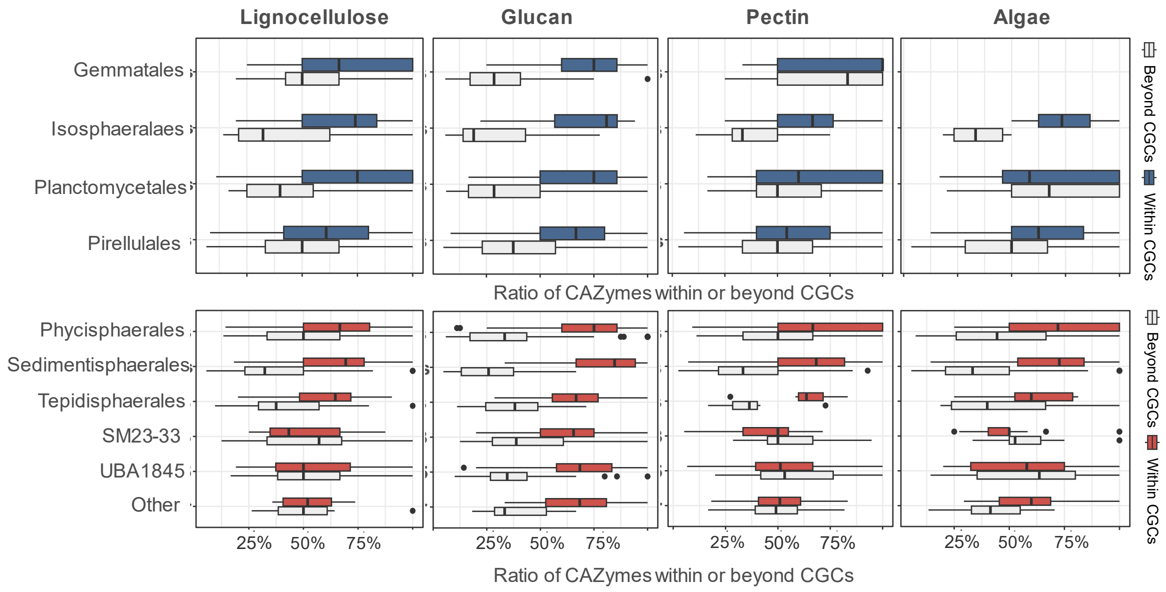


**Fig. S7.** CAZyme gene clustering of planctomycetotal CAZymes putatively engaged in the degradation of specific polysaccharides. Only orders of the Planctomycetia and Phycisphaerae classes are shown. GH and PL families within or beyond CGCs involved in the degradation of lignocellulose (cellulosic and hemicellulosic fractions), glucans (α-glucans), pectins and algal-derived polysaccharides. In light grey CAZymes outside CGCs.

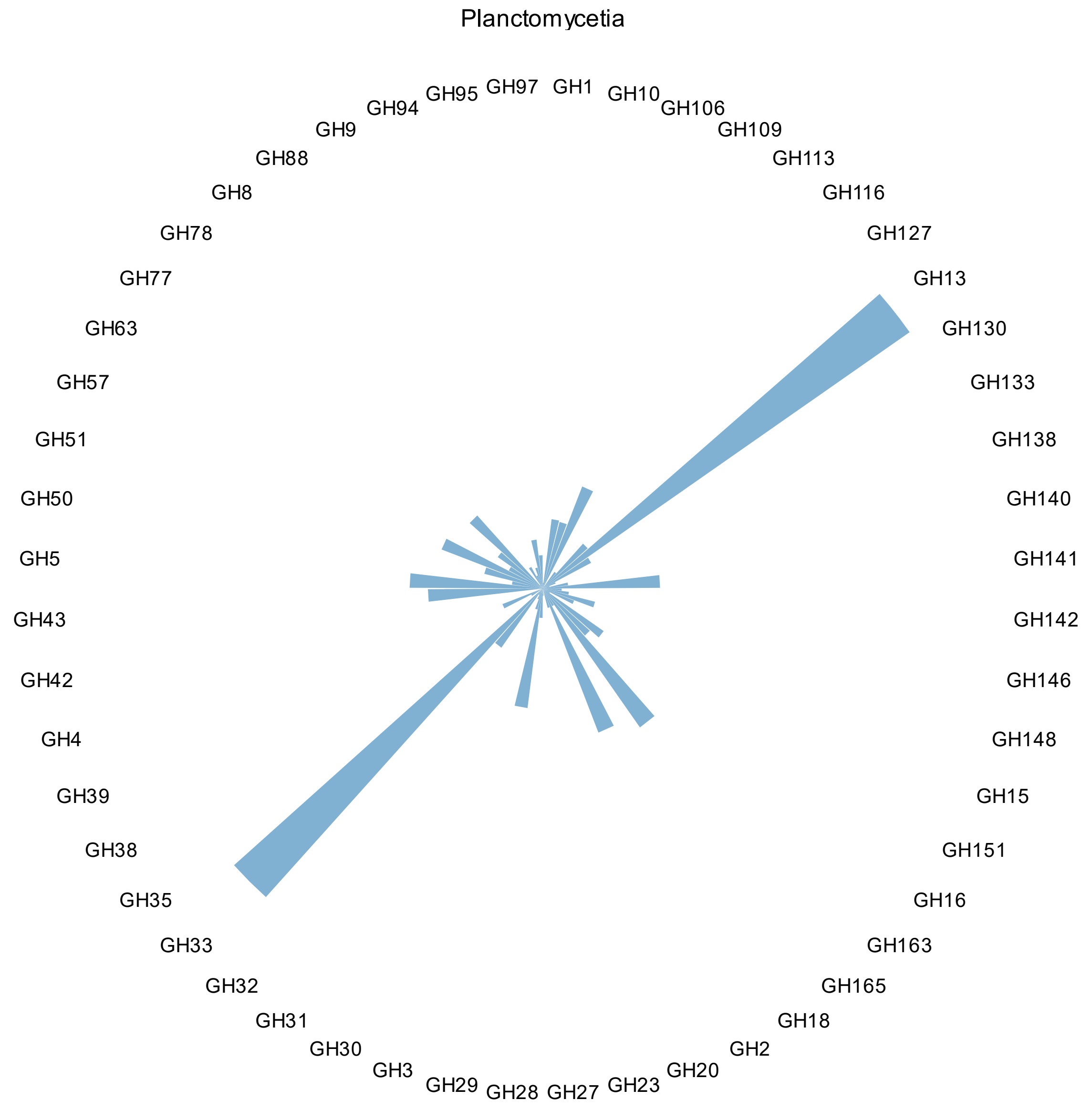


**Fig. S8.** Common co-localised CAZymes within CGCs for Planctomycetia class.

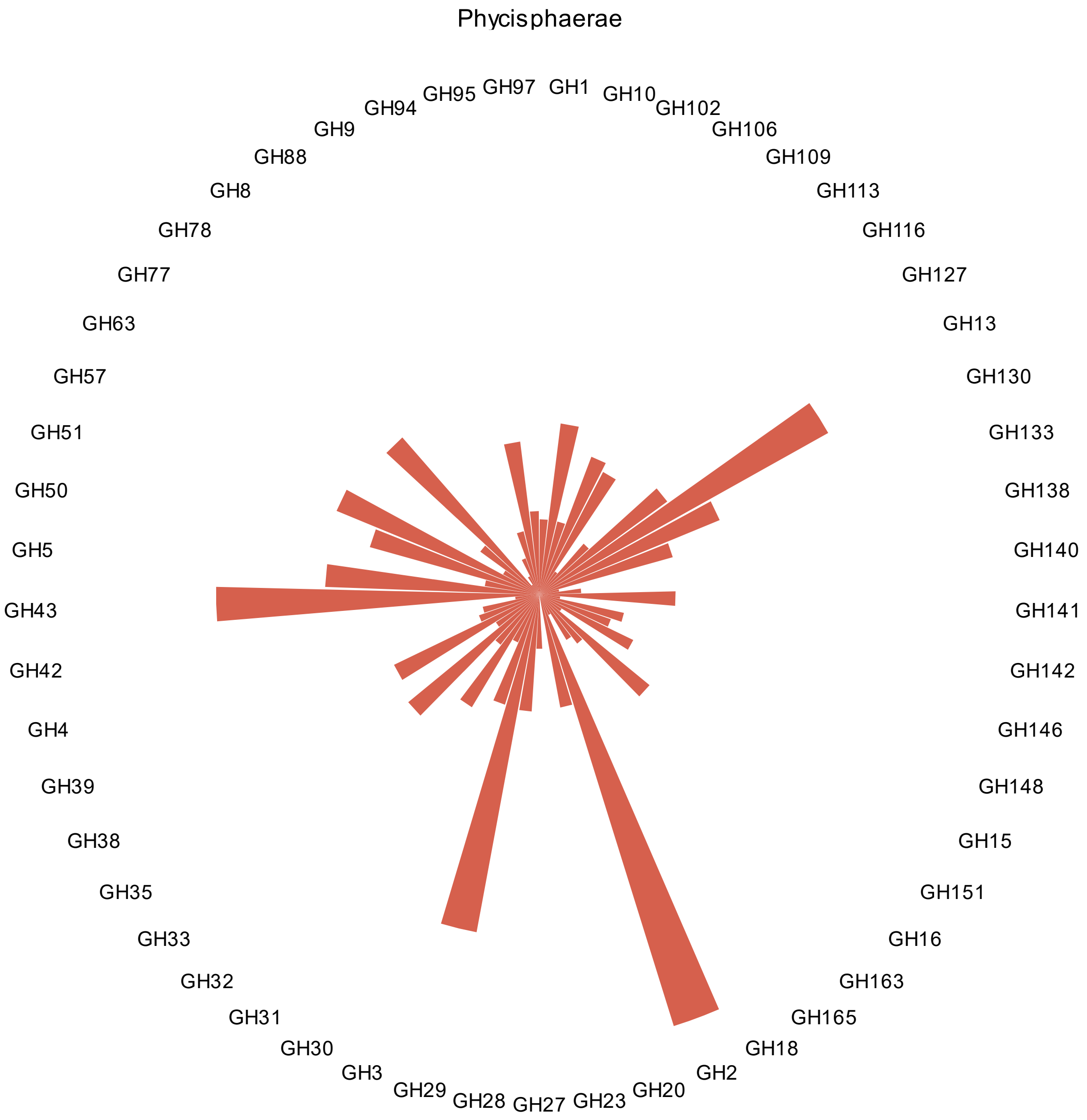


**Fig. S9.** Common co-localised CAZymes within CGCs for Phycisphaerae class.

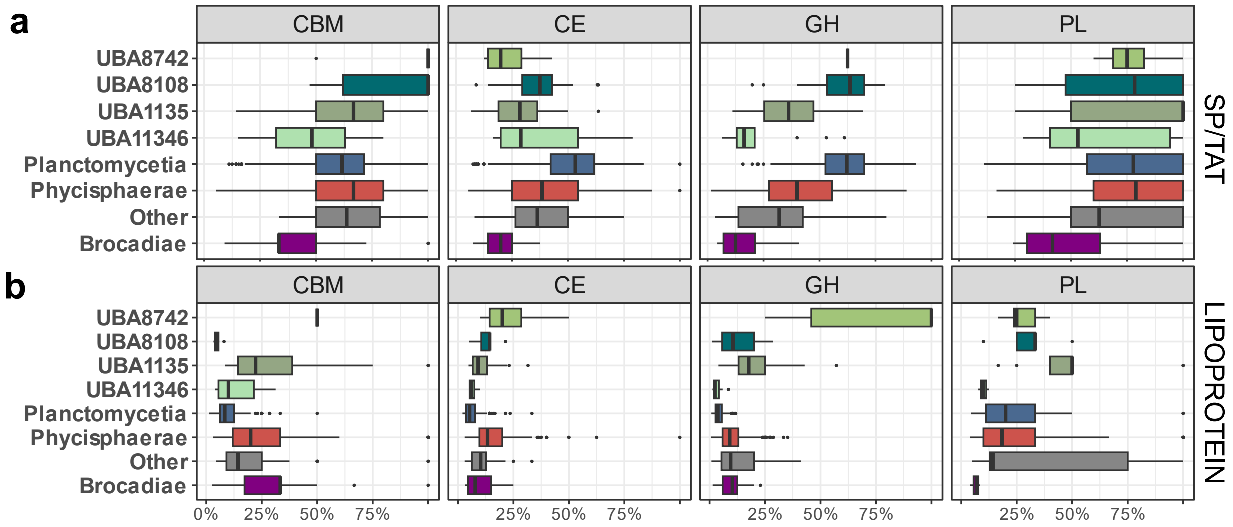


**Fig. S10.** Predicted localisation of CAZymes, coloured at the class level. Only main trends are shown (a) Ratio of CAZymes with predicted SP or TAT signal peptide for each planctomycetotal genome, grouped at the class level (b) Ratio of CAZymes with predicted signal peptides for lipoproteins for each planctomycetotal genome, grouped at the class level.
