## Additional File 8 for "Comparative genomic analysis of *Planctomycetota* potential towards complex polysaccharide degradation identifies phylogenetically distinct groups of biotechnologically relevant microbes"

### Supplementary Discussion

#### High potential for the discovery of novel and unique CAZymes in *Planctomycetota*

At present, planctomycetotal CAZymes remain largely unexplored and to prospect new functionalities in planctomycetotal genomes. Therefore, we evaluated the novelty of CAZymes by comparing their sequences to the entries in the CAZy database [4]. Overall, CAZymes encoded in *Planctomycetota* are distantly related to other bacterial CAZymes with the protein sequence identity oscillating on average between 48% and 62% (Additional File 2, Table S13). In general, the similarity of *Planctomycetia* CBMs, CEs, GHs and PLs oscillate around two peaks, with a larger fraction of CAZymes displaying around 60% of protein identity to entries in the CAZy database and only a smaller fraction characterised with high sequence identity. Interestingly, *Brocadiae* seem to encode CAZymes, mainly GHs, with the highest sequence homology to known families (>60%), while most of the other planctomycetotal CAZymes show a peak around the value of 50%. On the contrary, the AAs of planctomycetial *Isosphaerale*s, which encode the largest repertoire of AAs among the analysed planctomycetotal genomes, were distantly related to the entries in the CAZy database (~40%), putatively highlighting new functionalities (Additional File 2, Table S9). All the above observations signify that the bacterial

CAZy database only partially represents the diversity of CAZymes encoded by *Planctomycetota,* highlighting the potential for the discovery of novel CAZymes and the further need for the biochemical characterisation of these putatively functionally distinct enzymes.

A relatively large number of planctomycetotal CAZymes showed very low sequence identity to any previously characterised enzyme, i.e. below 30%, and all of these CAZymes were retrieved from *Planctomycetia, Phycisphaerae* and UBA1135 genomes (Additional File 2, Table S13). Among the CBM modules, those with the lowest sequence identity are versatile CBM51 and CBM57, rhamnose-binding GH67, fucose-binding CBM47 and cellulose-binding CBM9 and CBM16. Below the set threshold we found only a single PL8 family, putatively involved in the breakdown of various polysaccharides such as xanthan, chondroitin sulphate, alginate, and CE15 which typically displays ligninolytic activity (CAZy database). Many of the families identified included highly polyspecific GHs like GH2, GH5 or GH43, characterised with a broad variety of activities. Typically, these enzymes act on different oligosaccharides, including cellulosic and hemicellulosic derivates (Additional File 3, Table S1). Other unique CAZymes are assigned to GH78, GH106, GH139, GH141, which are mainly involved in the decomposition of pectins and algal-derived polysaccharides. Many of these CAZymes, including putative α-L-rhamnosidases (families GH78 and GH106), were previously shown to occur by past lateral gene transfers in different bacteria [108], indicating that the same event might have led to the evolution of gene variants found exclusively in planctomycetotal genomes. A high degree of planctomycetotal CAZyme sequence conservation could indicate that an ancestral gene first appeared in a *Planctomycetota* ancestor. Further gene duplications and numerous horizontal gene transfer events (HGT) might have allowed the resulting genes to spread between *Planctomycetota* and other PVC, contributing to the high diversity and uniqueness of contemporary planctomycetotal CAZyomes. The high degree of novelty within planctomycetotal CAZymes is in line with a previous study analysing a large group of β-galactosidase homologues from planctomycetotal genomes, which highlighted the presence of multiple, poorly characterised CAZymes, almost exclusively present in the PVC superphylum and some *Bacteroidota* [109][1]. Another recent study described the diversity of α-l-arabinofuranosidase homologues (GH51) from subantarctic intertidal sediments in different bacteria including *Planctomycetota* [110].

Finally, we also evaluated the abundance of what we called unclassified GHs, PLs, and CEs, that is, enzymes which were assigned by DIAMOND as putative CAZymes, but not classified to any of the currently recognised CAZyme families using HMMER-based sequence similarity approach (see Methods). *Planctomycetota* typically encode between 1 and 5% of unclassified CAZymes in their genomes (Fig. 4c). Genomes affiliated to *Gemmatales* encode the highest ratio of unassigned CAZymes, constituting on average 5% of their CAZyomes, most likely due to their greater genomic sizes (Fig. 3c and d). Recently, an exploratory study of the capybara gut microbiome genetic potential focused on genes annotated as hypothetical CAZymes, led to a discovery of a new GH173 family of β-galactosidases, and a new CBM89 family involved in xylan binding [111]. Therefore, further investigation of unclassified CAZymes, and CAZymes with a low sequence homology to known proteins, shall, in the future, allow the discovery of novel planctomycetotal functionalities as highlighted in the past by Naumoff and Dedysh [108].

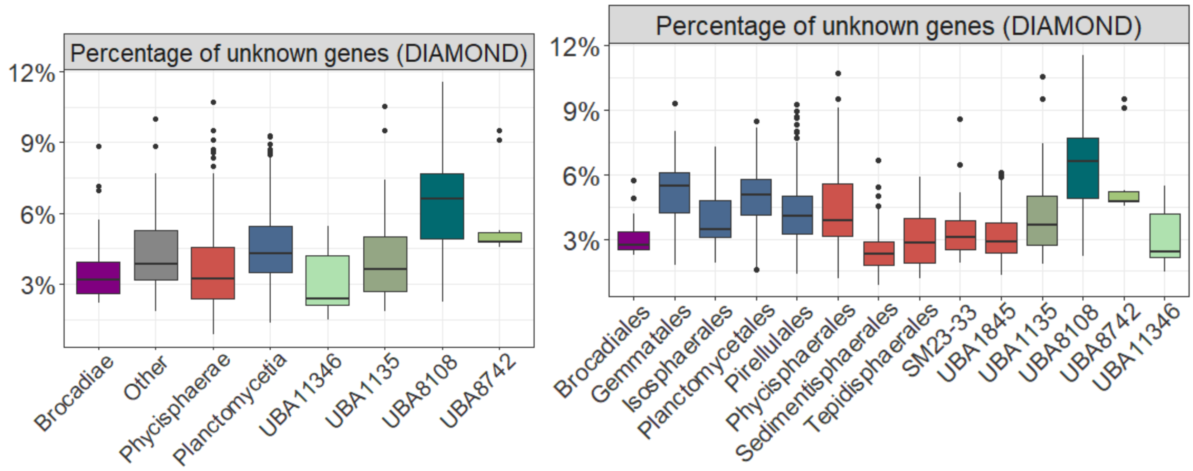

**Fig. S11.** The novelty and uniqueness of planctomycetotal CAZymes. Percentage of CAZymes within planctomycetotal genomes not assigned by DIAMOND to any known CAZyme family, grouped at class (a) and order (b) taxonomic level.

#### Lytic polysaccharide monooxygenases (LPMO) in genomes of *Planctomycetota*

Multiple studies have proven that the efficiency of lignocellulose (biomass) degradation can be remarkably improved by the joint action of lytic polysaccharide monooxygenases (LPMOs) and common GHs [58]. Interestingly, in the course of our analysis, we identified LPMO coding genes in many *Planctomycetota* genomes, mainly representing uncultured members of the *Phycisphaerae* class belonging to SM1A02 putative family, UBA8108 class and sporadically in other members of the phylum (Additional File 2, Table S3).

Planctomycetotal genomes regularly encode a single copy of AA10 gene, and based on protein phylogeny, three major clusters were observed that were separate from other, even phylogenetically unrelated bacteria representing different phyla such as *Actinomycetota, Pseudomonadota* and *Bacillota* (Fig. S12). This signifies that planctomycetotal AA10 enzymes are distantly related to all the other currently described LPMOs and accordingly, their sequence similarity to other publicly accessible proteins is assessed at between 28% and 68% (Additional File 2, Table S13). Furthermore, the sequence alignment of all the planctomycetotal AA10 proteins revealed only moderate coverage in a few regions (pairwise identity median of 25.8%), advocating for high intra-specialisation within this group (Additional File 2, Table S14). As such, the as yet uncharacterised planctomycetotal AA10 family might represent new hydrolytic functionalities that are not only distantly related to existing sequences in public databases, but also functionally diverse.

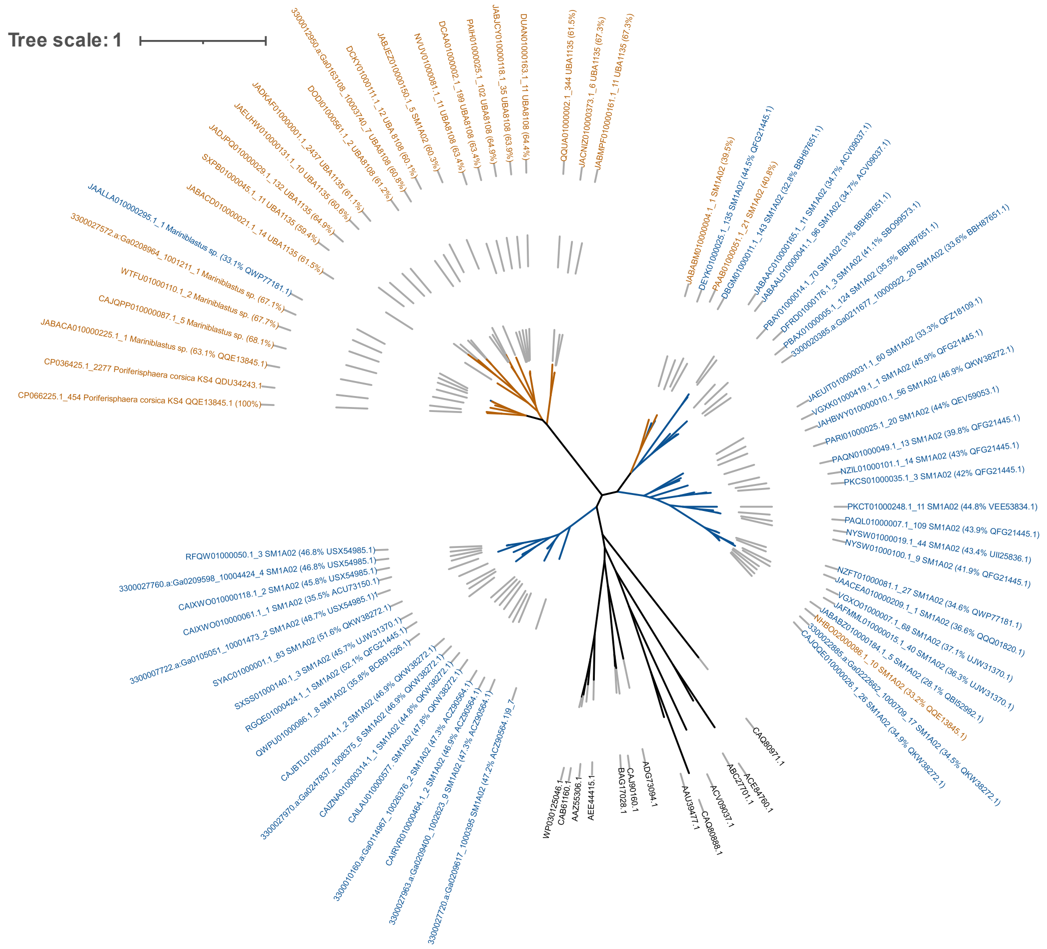


**Fig. S12.** Phylogeny of 57 planctomycetotal AA10 protein sequences (coloured in blue and brown) and 15 other AA10 enzymes with confirmed lytic oxidoreductase activity (coloured in black). Neighbor-Joining tree was built using aligned sequences by muscle algorithm. In brackets percentage identity of each protein and its closest blast hit (NCBI entry). Nodes coloured in brown represent proteins with the highest similarity to other *Planctomycetota* (*Poriferisphaera corsica* KS4*).* Nodes coloured in blue point to other donor bacteria such as *Actinomycetota, Bacillidota* and *Proteobacteroidota.* Nodes coloured in black represent enzymes already described in the literature.

In line with the predicted function of boosting GHs, most LPMOs bear either Sec or TAT signal peptide (>70%), indicating that they might be secreted. In addition, virtually half of the planctomycetotal LPMOs are multi-modular, and are appended to CBM2 (Fig. S13) which has an affinity to cellulose, and less often to chitin and xylan (CAZy database). Indeed, one multi-modular enzyme from a marine SM1A02 bacterium contains an additional GH18 module, which suggests its chitinolytic activity (Fig. S13). Overall, future investigations of CBM interactions with AA10s will be crucial for identifying the enzyme specificity but also for uncovering the process of the cellulosic and hemicellulosic polysaccharide breakdown in *Planctomycetota*.

LPMO coding genes are typically distributed beyond CAZyme clusters and only 20% of genomes were predicted to encode single CGCs containing AA10 (Supplementary Table 4, Table S2). Intriguingly, as opposed to freshwater bacterial CGCs, only genomes originated from freshwater metagenomes encoded in their operons other CAZymes besides AA10, including multi-domain enzymes. Among classified CAZyme domains we detected chitinases GH18, lignocellulose-targeting such as GH5, GH9 as well as CBM modules that bind either chitin CBM5, CBM12, CBM55, or cellulose CBM3, CBM49. Little is known about the carbohydrate active clusters involving LPMO genes, but for instance, some *Pseudoalteromonas* strains were identified with the operons encoding LPMO flanked with two GH18 chitinases [112]. We argue that LPMOs co-localised with other GHs might represent an evolutionary optimised version of an efficient enzymatic machinery, possibly targeting complex polysaccharides like crystalline cellulose or chitin. However, while LPMOs might specifically enhance the activity of co-localised glycosidases, LPMOs encoded beyond clusters might be universal boosters helping diverse enzymes to attack glycosidic bonds within the polysaccharide moieties. Cellular investment in a single enzyme production would offer an interesting cost-saving strategy compared to the expression of the whole enzyme cluster. At the same time, potent LPMOs would represent an interesting component of industrially relevant enzymatic preparations. The simplification of enzymatic cocktails would significantly reduce the cost of the enzymatic biomass processing technologies, e.g. for bioethanol production [59]. So far, none of the planctomycetotal LPMOs have ever been analysed, therefore their enzymatic activities should be further studied to determine their effectiveness for biomass.

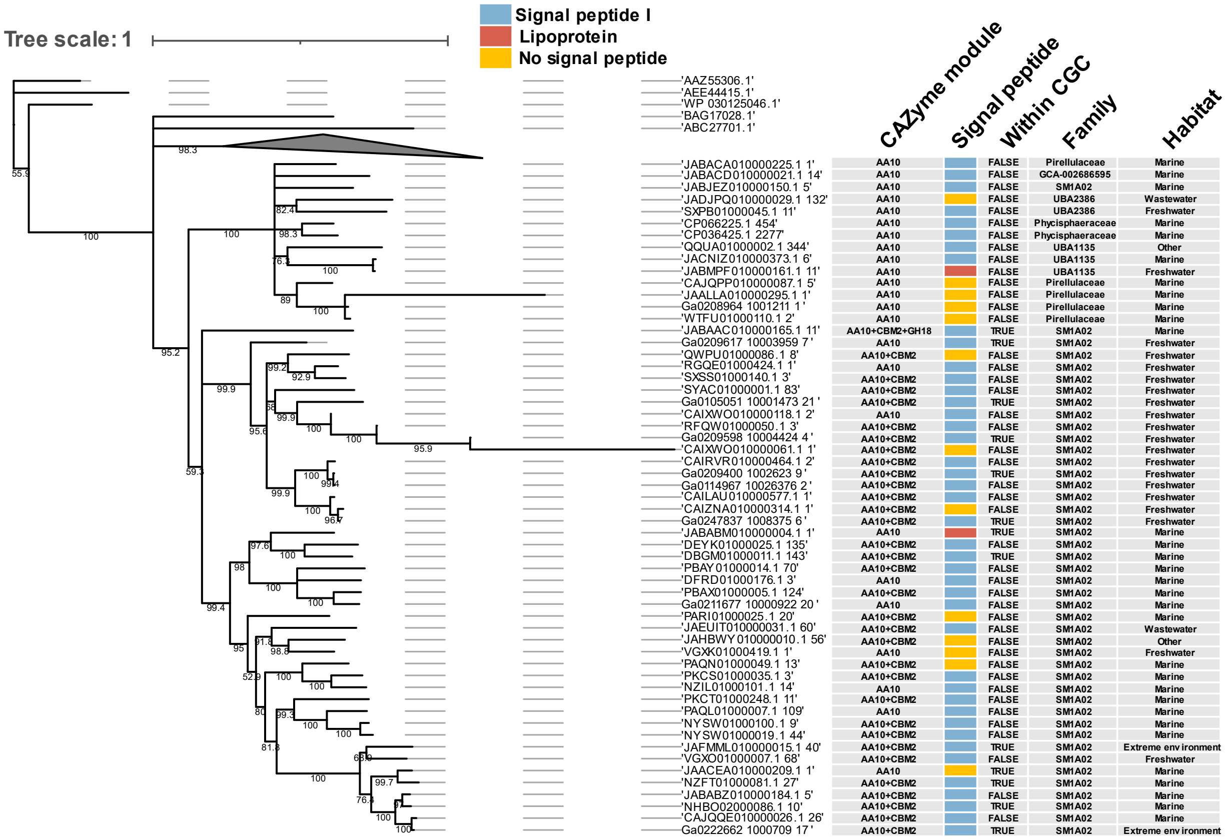


**Fig. S13.** The phylogeny of planctomycetotal AA10 protein sequences retrieved from genomes assigned to SM1A02 putative family and the additional 15 AA10 protein sequences from bacteria representing other phyla with the lytic polysaccharide monooxygenase activity described (LPMO). Bootstrap values are shown on branches.
